# Transposition, duplication, and divergence of the telomerase RNA underlies the evolution of *Mimulus* telomeres

**DOI:** 10.1101/2023.12.06.568249

**Authors:** Surbhi Kumawat, Irene Martinez, Dhenugen Logeswaran, Hongfei Chen, Jenn M. Coughlan, Julian J.-L. Chen, Yaowu Yuan, James M. Sobel, Jae Young Choi

## Abstract

Telomeres are nucleoprotein complexes with a crucial role of protecting chromosome ends. It consists of simple repeat sequences and dedicated telomere-binding proteins. Because of its vital functions, components of the telomere, for example its sequence, should be under strong evolutionary constraint. But across all plants, telomere sequences display a range of variation and the evolutionary mechanism driving this diversification is largely unknown. Here, we discovered in Monkeyflower (*Mimulus*) the telomere sequence is even variable between species. We investigated the basis of *Mimulus* telomere sequence evolution by studying the long noncoding telomerase RNA (TR), which is a core component of the telomere maintenance complex and determines the telomere sequence. We conducted total RNA-based *de novo* transcriptomics from 16 *Mimulus* species and analyzed reference genomes from 6 species, and discovered *Mimulus* species have evolved at least three different telomere sequences: (AAACCCT)*_n_*, (AAACCCG)*_n_*, and (AAACCG)*_n_*. Unexpectedly, we discovered several species with TR duplications and the paralogs had functional consequences that could influence telomere evolution. For instance, *M. lewisii* had two sequence-divergent TR paralogs and synthesized a telomere with sequence heterogeneity, consisting of AAACCG and AAACCCG repeats. Evolutionary analysis of the *M. lewisii* TR paralogs indicated it had arisen from a transposition-mediate duplication process. Further analysis of the TR from multiple *Mimulus* species showed the gene had frequently transposed and inserted into new chromosomal positions during *Mimulus* evolution. From our results, we propose the TR transposition, duplication, and divergence model to explain the evolutionary sequence turnovers in *Mimulus* and potentially all plant telomeres.

## Introduction

The ends of all linear eukaryotic chromosomes need special attention. Since DNA replication is a semiconservative process, without any intervention chromosome ends will progressively erode with each round of DNA replication (*i.e.* the end replication problem) (Watson 1972; Olovnikov 1973). In addition, naked chromosome ends trigger DNA damage responses in the cell and result in detrimental consequences, such as cellular senescence and chromosomal fusions (Shay and Wright 2005). To counteract these highly deleterious genome instabilities, chromosome ends are capped by a nucleoprotein complex called the telomere. The telomere is composed of a TG-rich microsatellite sequence (Podlevsky and Chen 2016) and the telomerase, a ribonucleoprotein enzyme complex, ensures the proper synthesis and maintenance of chromosome ends (Osterhage and Friedman 2009). Additional specialized telomere binding proteins also bind to the telomere and protect chromosome ends from being detected as damaged DNA (Vega et al. 2003; de Lange 2018).

All eukaryotes with a linear chromosome need the telomere for proper chromosome end protection (Fulcher et al. 2014). Due to this crucial function, telomeres are thought to be under strong evolutionary constraint and prohibited from molecular changes. For instance, all vertebrates have an identical telomere repeat motif TTAGGG that repeats multiple times at chromosome ends [we symbolize the telomere structure with a nucleotide sequence TTAGGG repeating *n* times at the chromosome end as (TTAGGG)*_n_*] (Meyne et al. 1989). It seems for vertebrates, extremely strong purifying selection has prohibited evolutionary changes in the telomere. But in plants, telomere sequences deviate from this ultra-constrained evolutionary history and display a diversity of telomere repeat sequences (Podlevsky and Chen 2016; Shakirov et al. 2022). *Arabidopsis thaliana* was the first plant species to have its telomere sequence decoded as (TTTAGGG)*_n_* (Richards and Ausubel 1988). Subsequent studies have then revealed many plant species also harbor the same *Arabidopsis*-type telomere sequence (Peska and Garcia 2020). Its common occurrence and similarity to animal telomere sequence (only a single nucleotide change) indicates (TTTAGGG)*_n_* is the ancestral telomere sequence in plants. But there are also multiple lineages deviating from the *Arabidopsis*-type sequence with sequence structures that can be grouped into three major types: (1) extension or contraction of the thymine or guanine nucleotide [*e.g.* (TTAGGG)*_n_* in the Asparagales group (Adams et al. 2001), (TTTTTTAGGG)*_n_* in *Cestrum elegans* (Peška et al. 2015), (AATGGGGGG)*_n_* in *Cyanidioschyzon merolae* (Nozaki et al. 2007)], (2) insertion or substitution of non-thymine or - guanine nucleotides [*e.g.* (TTCAGG)*_n_* and (TTTCAGG)*_n_* in the *Genlisea* group (Tran et al. 2015)], and (3) large changes with minimal resemblance to the *Arabidopsis*-type sequence [*e.g.* (CTCGGTTATGGG)*_n_* in the *Allium* group (Fajkus et al. 2016)].

The diversity of plant telomere sequences is fascinating and raises the question; what evolutionary processes are underlying this diversification? This question can be partially answered by studying the telomerase, which functions to maintain telomeres and synthesize telomeric DNA at chromosome ends. The telomerase is comprised of two major components (see Fig 1 for molecular model): 1) the telomere reverse transcriptase (TERT) that synthesizes the telomere DNA sequences and 2) the telomerase RNA (TR) a long noncoding RNA gene that acts as a template during the synthesis (Greider and Blackburn 1985; Greider and Blackburn 1989). Evolutionarily, TERT is found throughout the tree of life. In yeast, animal, and plant species with a telomerase-based mechanism of telomere maintenance, TERT is the main catalytic enzyme of the telomerase complex (Procházková Schrumpfová et al. 2019). Between species the length, sequence identity, and intron-exon structure of TERT varies considerably, but the core functional domains are largely conserved throughout the eukaryotic kingdom (Sýkorová and Fajkus 2009). The TR, on the other hand, is a rapidly evolving sequence, displaying extreme diversity in its size, sequence, and structure across eukaryotic clades (Podlevsky and Chen 2016). The level of sequence divergence is so elevated that often even between species from closely related genera, the TR sequence has minimal similarity that simple nucleotide BLAST based approaches have difficulties in determining orthology. Many long noncoding RNA genes are known to be rapidly evolving (Pang et al. 2006; Mattick et al. 2023) and the TR also shares this evolutionary pattern.

**Figure 1.**
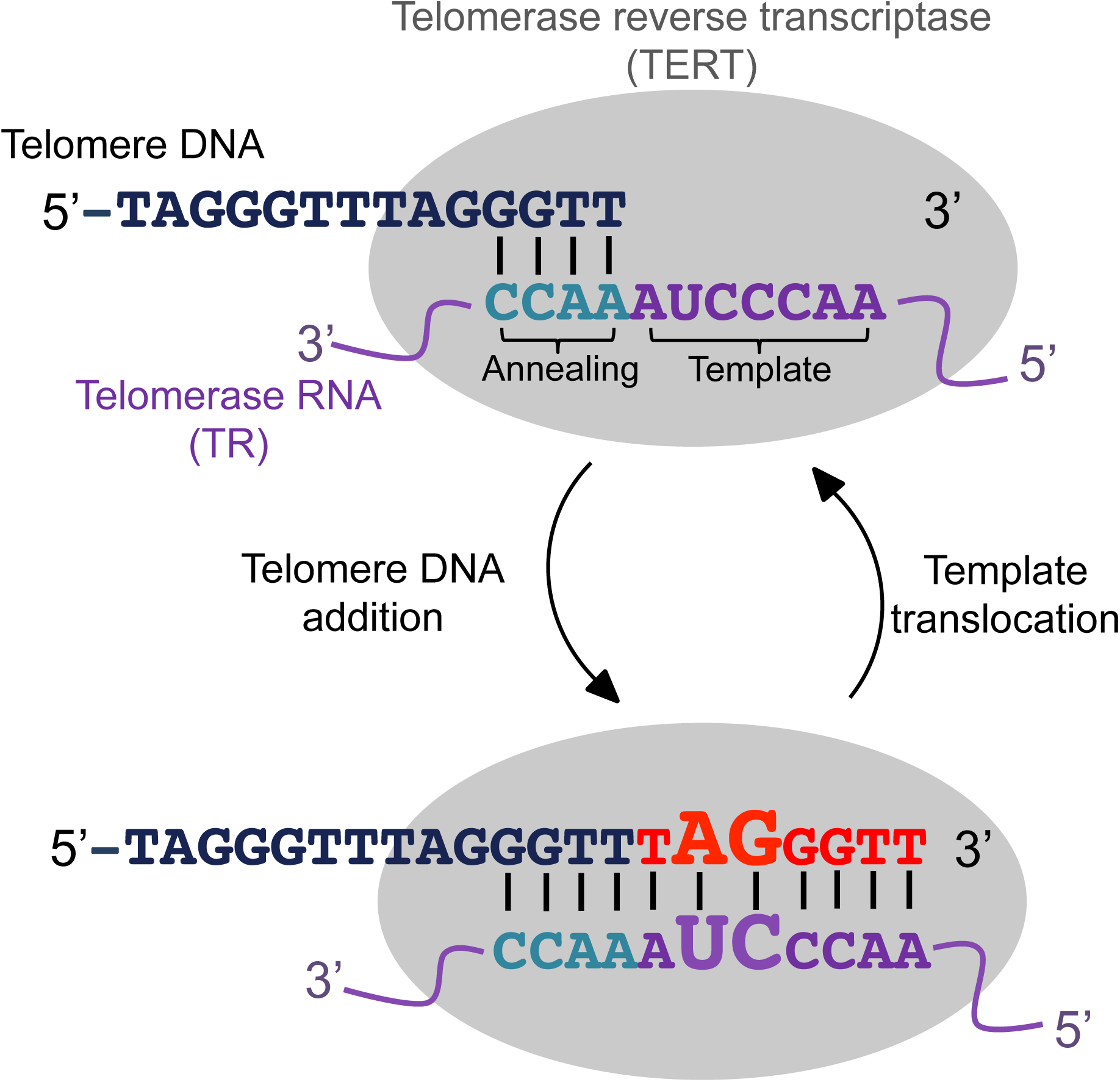
Hypothesized model for the *Mimulus* telomerase activity cycle. Figure was adapted from Kumawat and Choi (2023) and the activity model is based on Qi et al. (2012). The telomerase ribonucleoprotein complex comprises of TERT and TR, and functions to extend chromosomal ends with telomere DNA sequences. Within the telomerase catalytic domain the TR annealing sequence (teal letters) forms a DNA/RNA duplex with the 3’ ends of the telomeric DNA. TERT protein then synthesizes the deoxyribonucleotides to the 3’ end of the telomeric DNA (red letters) by using the TR templating sequence (purple letters) and through reverse transcription. After reaching the end of the templating sequence the DNA/RNA duplex separates and the TR templating domain reanneals with the telomeric DNA to open up the templating sequence for further synthesis of telomere DNA sequences. The variable telomere DNA sequence that we discovered in this study is highlighted as enlarged letters in the DNA/RNA duplex formed in the TR templating sequence region.

Across the sea of rapidly evolving sequences there are also “islands” of short stretches (1 to 8 bp) of highly conserved nucleotides that are scattered across the TR sequence (Podlevsky and Chen 2016). Plant TRs harbor 5 highly conserved regions (CRs) (Song et al. 2019) that are involved in folding the primary sequence into a higher order secondary structure resembling those found in animal TRs (Qi et al. 2013). In *A. thaliana* the CRs were functionally tested and shown to be crucial for proper telomerase activity (Song et al. 2019). In the 5’ region of the TR there are three conserved regions, CR1, CR2, and CR3 that folds into a template-pseudoknot (T-PK) domain and is involved in the catalytic activity of the telomerase (Podlevsky and Chen 2016). The remaining two domains (CR4 and CR5) in *A. thaliana* TR form a secondary structure with similarity to the helix IV domain in ciliates and CR4/5 domain in vertebrates. In ciliates, vertebrates, and plants this secondary structure has been shown to reconstitute the telomerase activity in *trans* with the T-PK domain (Mitchell and Collins 2000; Chen et al. 2002; Mason et al. 2003; Xie et al. 2008; Song et al. 2019). The CR1 domain, in particular, corresponds to the template domain and harbors two sequence motifs that are directly involved during the synthesis of the telomere DNA (see Fig 1 for summary of TERT-TR activity model). The first motif is the annealing sequence that binds to the single stranded telomere DNA and the second motif is the template sequence that serves as the template for TERT to synthesize telomeric DNA sequences (Greider 1991). Alignment between the telomere DNA and TR template domain is crucial for proper extension of the telomere and the plant species with divergent telomere sequences have also evolved the same changes in the TR template sequence (Fajkus et al. 2019; Song et al. 2019; Fajkus et al. 2021).

Studying how the TR has evolved in plants with divergent telomere sequences could uncover novel insights into the mechanism that underlies the evolution of plant telomere sequences. Currently, almost all previous plant telomere sequence studies have focused on the telomere sequences between species of divergent phylogenetic clades. While these comparisons have been greatly fruitful for surveying the telomeric sequence variation in plants, the deep evolutionary scale has prohibited from establishing the fine-scale evolutionary processes that are involved in the telomere sequence evolution. In this study we investigated the evolutionary basis of plant telomere sequence variation by studying the telomeres of the genus *Mimulus* sensu lato (Barker et al. 2012; Lowry et al. 2019). *Mimulus* belongs to the order Lamiales, which includes well known plants such as the classic genetic model Antirrhinum (family Plantaginaceae) and economically important crops like olive (family Oleaceae) and sesame (family Pedaliaceae) (Refulio-Rodriguez and Olmstead 2014). The genus consists of ∼170 species of herbaceous annual, herbaceous perennial, and woody perennial plants that are found in a wide range of habitats throughout the California Floristic Zone (Beardsley et al. 2004). In the past, *Mimulus* has been a popular system for conducting ecological and evolutionary biology research and has provided key fundamental insights into the genetics and ecology of adaptation and speciation (Wu et al. 2008; Yuan 2019). Its telomeres, on the other hand, have not been investigated before.

Here, we took a genomic approach and studied the evolution of the telomere in 18 *Mimulus* species, with a goal to understand the evolutionary basis of plant telomere sequence variation. We investigated the evolution of the *Mimulus* TR gene through *de novo* transcriptomics and annotating the TR gene in available reference genome assemblies for a deeper evolutionary understanding of the telomere sequence variation. Based on our results we propose the telomerase RNA transposition, duplication, and divergence model and hypothesize the turnover in plant telomere sequences is an evolutionary outcome from a transposition-mediated duplication of the TR gene. The duplicate TR opens up the evolutionary opportunities for sequence divergence in the templating sequence of the TR, resulting in a telomere with sequence heterogeneity. Our model provides a novel evolutionary mechanism to explain the telomere sequence turnover in *Mimulus* and potentially for all plants.

## Result and Discussion

### K-mer analysis on genome sequencing data uncovers the telomere sequence variation between *Mimulus* species

This project was originally motivated by an attempt to characterize the telomere length variation across *Mimulus* species. In our previous study (Choi et al. 2021), we used the computational method k-Seek (Wei et al. 2014; Wei et al. 2018) to count k-mer repeats in raw un-mapped whole genome sequencing reads, and used the telomere repeat k-mer counts as an approximation of the telomere length. We previously showed this computational approach had high correlation with direct telomere length measurements (ρ = 0.55) and was a reliable *in silico* method to quantify the telomeric repeats in the genome. We used k-Seek to analyze published whole-genome re-sequencing data of *Mimulus* species from three different sections (*i.e.* morphological groupings) and our original aim was to analyze the telomere length variation between species of divergent *Mimulus* section (family Phrymaceae). The species we examined the genome sequencing data were *M. aurantiacus* from section *Diplacus* (Stankowski et al. 2019), *M. guttatus* from section *Simiolus* (Troth et al. 2018), and *M. verbenaceus* from section *Erythranthe* (LaFountain et al. 2023). The section *Simiolus* and *Erythranthe* are phylogenetic sisters while *Diplacus* is the third outgroup (Beardsley and Olmstead 2002) and the three sections last shared a common ancestor 32–50 million years ago (Kumar et al. 2017). Recent phylogenetic analyses indicate polyphyly of the genus *Mimulus* (MoralesLBriones et al. 2022), and taxonomic revision has split these focal species into separate genera, *Diplacus* and *Erythranthe* (Barker et al. 2012). However, following other recent work in the Phrymaceae, we have elected to retain the use of the name *Mimulus* for its historic significance and recognizability (see Lowry et al. 2019).

We used our published k-Seek analysis pipeline (Choi et al. 2021) to characterize the genome-wide k-mer abundance across the three focal *Mimulus* species (*M. aurantiacus*, *M. guttatus*, and *M. verbenaceus*). By analyzing the population genomic data we were able to quantify the natural variation in k-mer counts across multiple genotypes for each species (n = 43 for *M. aurantiacus*, n = 228 for *M. guttatus*, and n = 54 for *M. verbenaceus*). Note for *M. aurantiacus*, the population genomic data are from six closely related subspecies that we are collectively referring to as *M. aurantiacus* only for the k-mer analysis for convenience, and importantly our k-mer results do not depend on the underlying subspecies classification. Results showed for all three species, most large sized k-mers (k > 3) had relatively low abundance (∼100> copies per 1× genome coverage), indicating the genome-wide tandem repeat variation was largely driven by small sized 1-, 2-, or 3-mers (Supplementary Fig. S1). But in *M. aurantiacus* (Fig. 1), the k-mer AAACCCT had substantial abundance for a large sized k-mer (13,190 median copies per 1× genome coverage) and the sequence corresponded to the *Arabidopsis*-type telomere repeat sequence (note AAACCCT is a offset of the reverse complement of TTTAGGG [*i.e.* CCCTAAA] followed by tandem repetition). We then took an independent dataset to check whether the AAACCCT k-mer corresponded to the telomere repeat sequence. We analyzed the *M. aurantiacus* reference genome assembly, which was a chromosome-level genome assembly and sequenced from the subspecies puniceus (Stankowski et al. 2019), and focused on the sequences at the ends of assembled chromosome sequences to identify the telomere repeat motif. *M. aurantiacus* ssp. puniceus The chromosome ends were abundant for the k-mer AAACCCT (Supplementary Fig. S2) indicating *M. aurantiacus* ssp. puniceus had the canonical *Arabidopsis*-type telomere sequence.

The genome sequencing k-mer analysis for *M. guttatus* and *M. verbenaceus*, on the other hand, did not detect AAACCCT as the most abundant 7-mer repeat (Fig. 2). In fact AAACCCT was almost absent in *M. guttatus* (0 median copies per 1× genome coverage) while it had a very low abundance in *M. verbenaceus* (147 median copies per 1× genome coverage). This suggested the telomere sequence motif had changed in *M. guttatus* and *M. verbenaceus*. To investigate the alternative telomere motif we searched for k-mers that resembled the *Arabidopsis*-type telomere sequence and had high abundance in the raw genome sequencing data. In *M. guttatus* the k-mer AAACCCG was the highest abundant 7-mer sequence (645 median copies per 1× genome coverage), and this k-mer was largely absent in *M. aurantiacus* (10 median copies per 1× genome coverage) and *M. verbenaceus* (0 median copies per 1× genome coverage). Meanwhile in *M. verbenaceus* there was no highly abundant 7-mer sequence but the 6-mer AAACCG was highly abundant (6,070 median copies per 1× genome coverage), and this k-mer was largely absent in *M. aurantiacus* (0 median copies per 1× genome coverage) and *M. guttatus* (7 median copies per 1× genome coverage). To investigate if these two k-mers corresponded to the telomere repeat we analyzed the chromosome-level reference genome assemblies for *M. guttatus* and *M. verbenaceus*. We focused on the sequences at the chromosome ends and discovered *M. guttatus* were enriched for the AAACCCG repeat and *M. verbenaceus* were enriched for the AAACCG repeat (Supplementary Fig. S2), which indicated the abundant k-mers that resembled a telomere repeat from the *M. guttatus* and *M. verbenaceus* raw genome sequencing data (Fig. 2) were likely to correspond to the telomere sequence.

**Figure 2.**
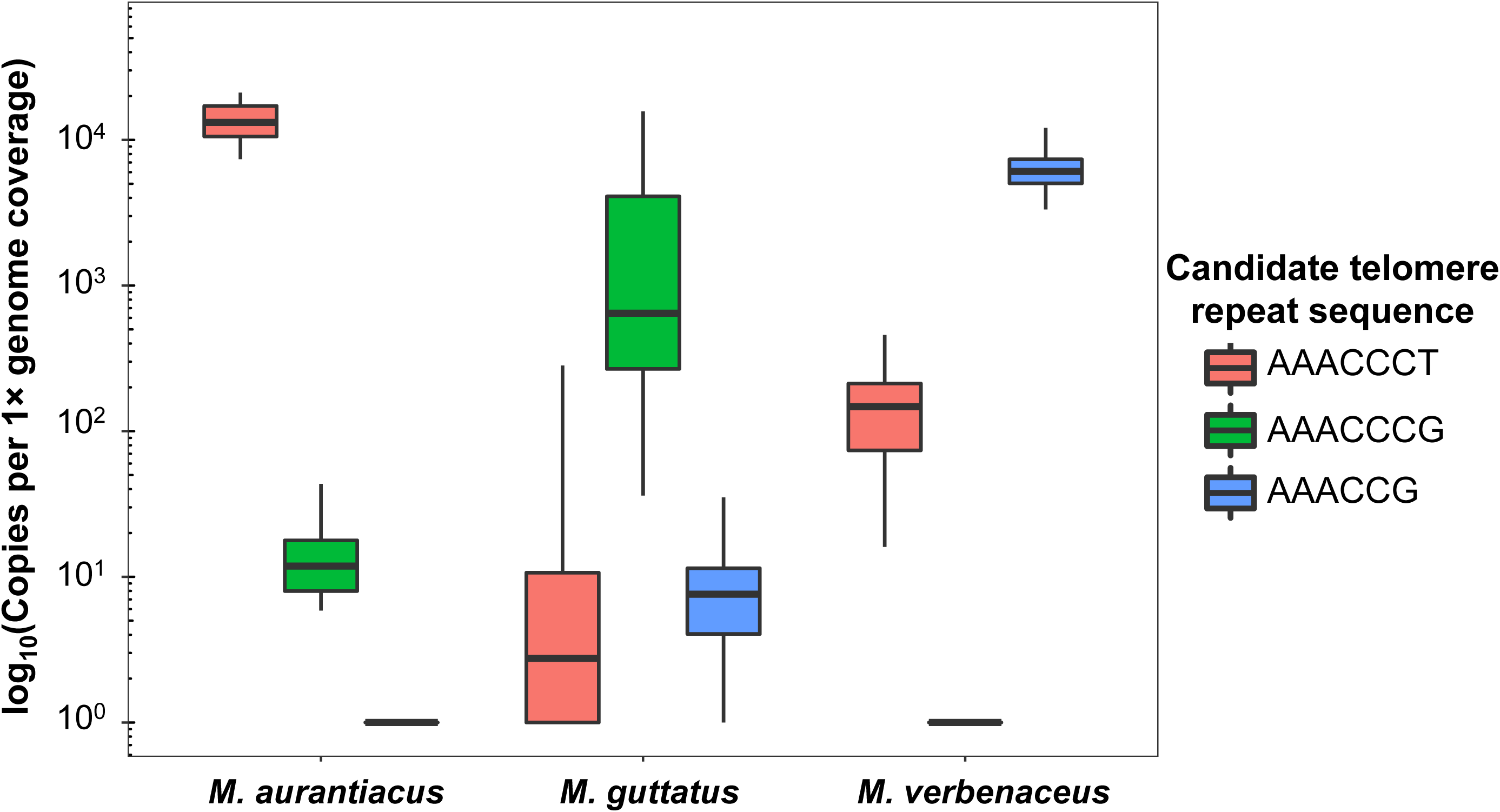
K-mer based genome sequencing analysis shows changes in the telomere sequence for three *Mimulus* species. For each species the abundance of the top three k-mers that resemble a telomere sequence are only shown. Within a species the most abundant k-mer is hypothesized as the species telomere sequence, while the other two k-mers are likely to be low abundance intergenic repeats.

### Annotating the Telomerase RNA gene in the *Mimulus* species with reference genomes

Our k-mer analysis suggested the telomere sequence had changed at least twice in the *Mimulus* genus (*i.e.* AAACCCT, AAACCCG, AAACCG). To further investigate this possibility we studied the evolution of the *Mimulus* TR gene. The template sequence within the TR determines the telomere DNA sequence (Fig. 1), hence by investigating the evolution of the TR gene we aimed to gain a deeper understanding of the telomere sequence evolution in the *Mimulus* genus. We first analyzed the *Mimulus* species with reference genomes (*M. aurantiacus* ssp. puniceus, *M. cardinalis*, *M. guttatus*, *M. lewisii, M. parishii*, and *M. verbenaceus*) to identify the TR gene. *Mimulus cardinalis*, *M. lewisii,* and *M. parishii* are species that were not analyzed in our k-mer analysis and belongs to the *Erythranthe* section, which includes *M. verbenaceus* that was analyzed in our k-mer analysis (Fig. 2). For each reference genome, we used a position weight matrix (Mosig et al. 2007) and secondary structure-based algorithm (Nawrocki and Eddy 2013) to search and annotate the TR genes locations (see Table 1 for genome coordinates). The size of the TR ranged between 278 bp to 317 bp, which are similar in size to other TRs found across the plant kingdom (ranging from 234–390 bp) (Song et al. 2019; Fajkus et al. 2021). During the TR annotation we found an intriguing result where the reference genomes from *M. aurantiacus* ssp. puniceus, *M. cardinalis*, and *M. lewisii* had evidence of two TR gene annotations. The evolution of these TR duplicates will be further discussed in a later section of this study.

**Table 1.** Genome coordinate and sense of the telomerase RNA (TR) gene in species with a reference genome. For each TR gene its syntenic region in other species is also shown. The syntenic region was determined from orthology of the gene directly upstream and downstream of the TR gene. The reported coordinates in the syntenic region are the end position of the upstream gene and start position of the downstream gene. Note because of micro-synteny differences the exact location of a TR and its syntenic position may not completely overlap (*e.g.* see TR2 gene position in *M. lewisii* reference genome and the syntenic position based on *M. cardinalis* TR2).

To examine the potential functionality of the annotated TRs we extracted 100 bp upstream of the predicted start site of the TR transcript to look for conserved regulatory elements. Previous studies have found the upstream genetic regions of land plant TRs contained a highly conserved type 3 promoter motif called the Upstream Sequence Element (USE) and a TATA box motif (Fajkus et al. 2019; Fajkus et al. 2021). All of our candidate TRs, including the duplicates, had a USE motif TCCCACAT within the upstream region (Supplementary Fig. S3). This sequence was identical to the USE motif found in all surveyed Lamiales species. One exception was for *M. cardinalis* where one of its TR duplicates had a single nucleotide substitution in the conserved USE motif (TCCCAC**G**T; the nucleotide difference is bolded) and we verified this nucleotide change with Sanger sequencing. For all TR candidate sequences we also discovered the TATA box 28–30 bp downstream from the USE motif (Supplementary Fig. S3). In sum, the presence of highly conserved transcription initiation sequences indicated our annotated TRs were likely to be expressed.

### Identifying the Telomerase RNA through total RNA transcriptomes of multiple *Mimulus* species

We aimed to obtain the TR sequence from multiple *Mimulus* species through *de novo* transcriptomics and assemble the TR transcript from the total RNA pool. But first to understand which tissue the TR would most likely be expressed in, we conducted RT-PCR experiments to amplify the TR gene in several tissues across multiple *Mimulus* species. We used the TR sequence that was annotated from the *Mimulus* species with a reference genome to design primers and conducted RT-PCR analysis on cDNA generated from total RNA extracted from mature leaf, root, and floral meristem tissues. We chose the three divergent *Mimulus* species (*M. aurantiacus* ssp. puniceus, *M. guttatus*, and *M. verbenaceus*) that we initially conducted the k-mer and reference genome analysis (Fig. 2 and Supplementary Fig. S2) and conducted the RT-PCR analysis. Prior studies have discovered high expression of the TR in plant tissues with actively dividing cells (Fajkus et al. 2019; Song et al. 2019) and we also detected expression of the TR gene in the floral meristem for all three *Mimulus* species (Fig. 3A and see Supplementary Fig. S4 for negative PCR results when raw total RNA was used for RT-PCR). There was also evidence of expression in other tissues that are largely composed of differentiated cells (root and mature leaf).

**Figure 3.**
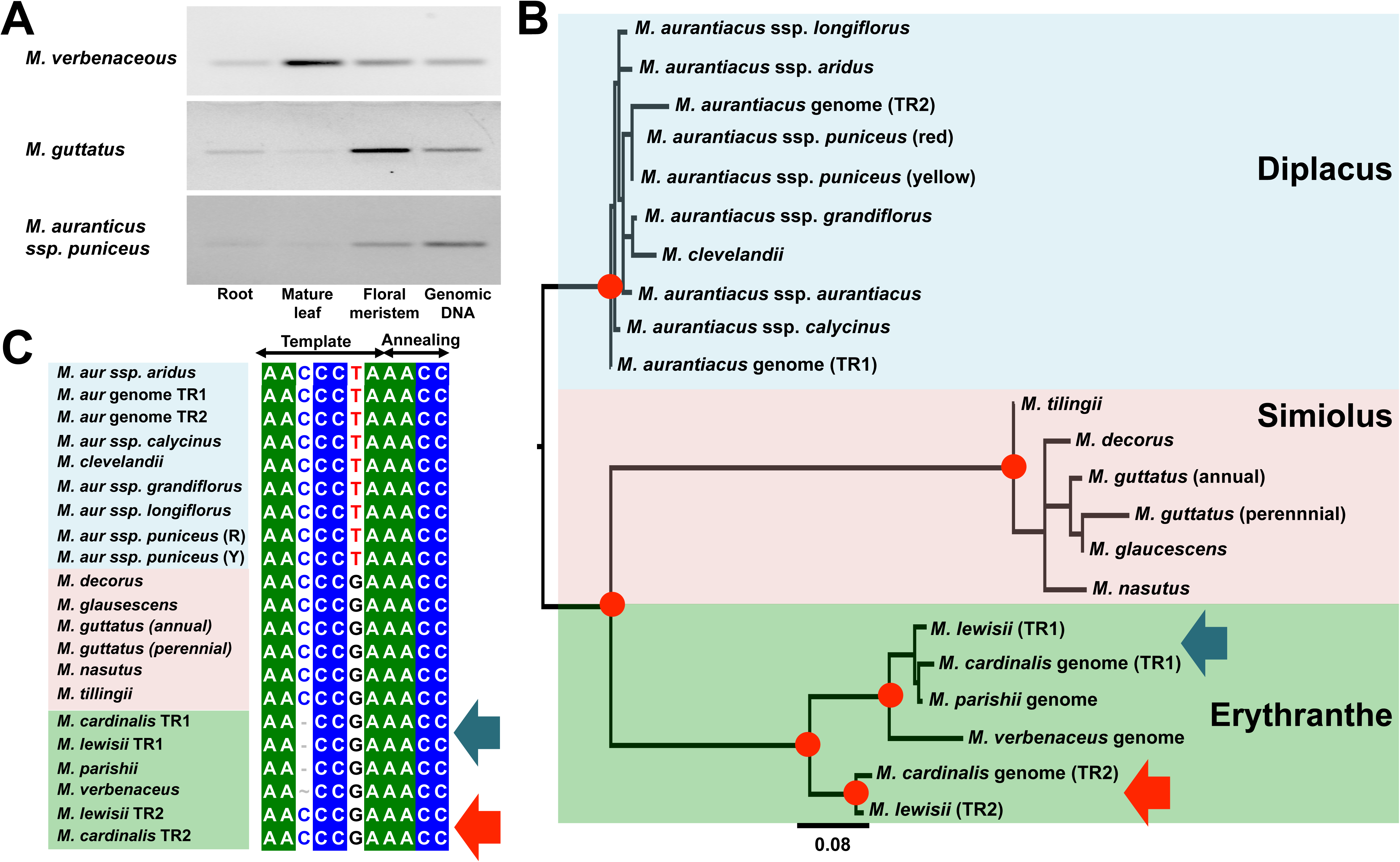
Molecular evolutionary analysis of the *Mimulus* Telomerase RNA (TR) gene. (A) RT-PCR results amplifying TR transcripts on cDNA generated from RNA extractions of three tissues (root, mature leaf, and floral meristem). Genomic DNA is shown as a positive control. (B) Phylogeny of the entire TR sequences assembled from 18 *Mimulus* species/subspecies. Sequences were obtained from total RNA transcriptome sequencing or by annotating the TR sequences from reference genomes (*i.e. M. aurantiacus* ssp. puniceus, *M. cardinalis*, *M. parishii*, and *M. verbenaceus*). Nodes with bootstrap support >95% are indicated with a red circle. Phylogenetic groups are colored according to the three *Mimulus* sections. Arrows point to the *M. cardinalis* and *M. lewisii* TR duplication. Blue arrow indicates TR1 and the red arrow indicates TR2. (C) Alignment of the TR templating region (see Figure 1 for involvement of sequence in the telomerase complex during telomere DNA synthesis). Sequences have been highlighted according to the color scheme of (B). Note based on the templating sequence, the *Diplacus* section will synthesize a AAACCCT telomere and *Simiolus* section a AAACCCG telomere. In *Erythranthe* section, all species will synthesize a AAACCG telomere, meanwhile in *M. cardinalis* and *M. lewisii* because TR2 has a AAACCCG templating sequence both species could have a potentially sequence heterogeneous telomere structure (but see Fig. 4 for results on *M. cardinalis* and *M. lewisii* TR duplication on telomere sequence).

Based on the RT-PCR results we targeted the floral meristem for total RNA transcriptome sequencing with a goal to assemble the TR sequence in multiple *Mimulus* species. We examined the floral meristem transcriptomes from 16 *Mimulus* species/subspecies and these include 8 (sub)species from the *Diplacus* section (*M. aurantiacus* ssp. *aridus*, *M. aurantiacus* ssp. *aurantiacus*, *M. aurantiacus* ssp. *calycinus*, *M. clevelandii*, *M. aurantiacus* ssp. *grandiflorus*, *M. aurantiacus* ssp. *longiflorus*, red flower ecotype *M. aurantiacus* ssp. *puniceus*, and yellow flower ecotype *aurantiacus* ssp. *M. puniceus*), 7 (sub)species from the *Simiolus* section (*M. decorus*, *M. glaucescens*, coastal perennial *M. guttatus*, inland perennial *M. guttatus*, inland annual *M. guttatus*, *M. nasutus,* and *M. tilingii*), and 1 species from the *Erythranthe* section (*M. lewisii*). We also took the TR sequences annotated from the reference genome (listed in Table 1) for a combined total of 19 (sub)species (see Supplementary Table 1 for list of analyzed species) TR sequences were available for downstream analysis.

The total number of raw reads generated for each transcriptome ranged from 102–182 million reads and more than 98% of those reads were recovered after quality control trimming to remove adapters and low quality sequence regions (Supplementary Table 2). We then aligned the quality control trimmed reads back to a reference genome to check for the proportion of reads that originated from a genome. Using *M. aurantiacus* ssp. puniceus reference genome for *Diplacus* species transcriptomes, *M. guttatus* reference genome for *Simiolus* species transcriptomes, and *M. lewisii* reference genome for the *M. lewisii* transcriptome, we determined there were between 50.0–99.7% of the reads aligned back to a reference genome. The low alignment percentage for some species is likely due to the reference genome being too divergent from the species with RNA-seq reads to be mapped.

Using the quality control trimmed sequencing reads, we used the trinity pipeline (Grabherr et al. 2011) to conduct *de novo* transcriptome assembly. After trinity based assembly the contig N50 ranged from 617–2619 bp and the number of assembled transcripts ranged from 117,776–266,437 (Supplementary Table 2). BUSCO scores were calculated and ranged from 26.4–70.4%. These low BUSCO scores may be related to our total RNA based sequencing strategy. We depleted rRNA but omitted a polyadenylated transcript selection step to enrich noncoding RNA sequences such as the TR, but this does not control for the vastly over abundant organellar (chloroplast and mitochondria) transcripts that are not polyadenylated (Forsythe et al. 2022). Consequently nuclear gene transcripts are likely to be undersampled, potentially resulting in the suboptimal assembly for coding sequences. In addition, because we sequenced the total RNA, the mRNA pool is also enriched for precursor messenger RNA (pre-mRNA) that did not undergo post-translational modification (Kukurba and Montgomery 2015). Here, the transcript assemblies will contain a mix of coding and noncoding sequences that will not be ideal for BUSCO as it uses open reading frames (ORFs) as a criterion for the transcriptome analysis (Simão et al. 2015). Despite the limitations we took the *de novo* transcriptome assembly and using the TR gene annotations from the reference genome assemblies, a BLAST-based nucleotide similarity analysis was able to identify the TR from 15 of the sequenced transcriptomes (no TR was successfully identified for the inland perennial *M. guttatus*). In the end, we were able to obtain the TR sequence from 15 *Mimulus* species (*M. aurantiacus* ssp. *aridus*, *M. aurantiacus* ssp. *aurantiacus*, *M. aurantiacus* ssp. *calycinus*, *M. clevelandii*, *M. aurantiacus* ssp. *grandiflorus*, *M. aurantiacus* ssp. *longiflorus*, red flower ecotype *M. aurantiacus* ssp. *puniceus*, yellow flower ecotype *aurantiacus* ssp. *M. puniceus, M. decorus*, *M. glaucescens*, coastal perennial *M. guttatus*, inland annual *M. guttatus*, *M. nasutus, M. tilingii*, and *M. lewisii*) from the transcriptome data and an additional TR sequences from 3 *Mimulus* species annotated from the genome assemblies (*M. cardinalis*, *M. parishii*, and *M. verbenaceus*) for a total of 18 unique *Mimulus* species.

### Genomic and transcriptomic data uncovers evidence of gene duplication for the telomerase RNA in a subset of *Mimulus* species

The *Mimulus* TR sequences were aligned to each other with MUSCLE (Edgar 2004) and the multi-sequence alignment was used to build a maximum-likelihood based phylogenetic tree to infer the evolutionary relationships (Fig 3B). The tree had high bootstrap support on internal nodes that grouped the species by *Mimulus* section (>99% after 1,000 bootstrap replicates) and the internal branch lengths between sections were deep. This phylogenetic evidence indicated there were substantial nucleotide differences between the TR sequences for species from different sections. This was also visually apparent in the multi-sequence alignment of the TR gene (Supplementary Fig. S5), which displayed a considerable amount of insertions and deletions (indels) of variable sizes between *Mimulus* sections.

During the genome annotation and transcriptome-based analysis of the TR gene, we discovered a surprising result where the TR gene was duplicated in several *Mimulus* species (*M. aurantiacus* ssp. puniceus, *M. cardinalis*, and *M. lewisii*). This was unexpected since TERT binds with a single TR molecule to extend chromosomal ends (Fig. 1), hence it was not immediately apparent why certain species had evolved a duplication in the TR gene. We then focused our analysis on the *Mimulus* TR paralogs to gain a deeper evolutionary understanding of the duplication event.

In *M. aurantiacus* ssp. puniceus, the reference genome contained two TR gene copies, but from our *de novo* assembly each *Diplacus* section species and subspecies had a single TR gene assembled. We randomly assigned the two TR copies from the *M. aurantiacus* ssp. puniceus reference genome as TR1 and TR2. Across the 244 aligned basepairs between *M. aurantiacus* ssp. puniceus TR1 and TR2 sequences, there were 11 nucleotide differences and a 50 bp insertion found only in TR2 (Supplementary Fig. S5). We investigated the phylogenetic relationship between *M. aurantiacus* ssp. puniceus TR1 and TR2 from the reference genome, and the rest of the *Diplacus* section TR, but the bootstrap supports were too low to make any confident evolutionary inferences (Fig. 3B). This suggested a recent evolutionary origin for the TR duplication in *M. aurantiacus* ssp. puniceus. In addition, we sequenced the transcriptomes of two *M. aurantiacus* ssp. puniceus ecotypes and in both subspecies we were only able to assemble one TR sequence. This indicated between the TR1 and TR2 paralogs annotated in the *M. aurantiacus* ssp. puniceus reference genome only one paralog was potentially expressed. In our RT-PCR results (Fig. 3A) we only detected a single band and given that the *M. aurantiacus* ssp. puniceus TR1 and TR2 from the reference genome had a 50 bp difference, if both genes were expressed we would expect to see two bands but the single TR band further indicated that only one of the TR paralogs were being expressed.

Within the *Erythranthe* section we annotated two TR copies for *M. cardinalis* and *M. lewisii* using their reference genome assembly. With the *M. lewisii* total RNA transcriptome, we were able to *de novo* assemble both TR genes, indicating the two TR paralogs were being expressed in the *M. lewisii* floral meristem that was extracted for the transcriptomics. There was strong phylogenetic support (>95% after 1,000 bootstrap replicates) that grouped TR by sequence and not by species. We designated the TR paralog that grouped with *M. parishii*, and *M. verbenaceus* as TR1, while the duplicate only found in *M. cardinalis* and *M. lewisii* as TR2. Since TR1 and its orthologs were found in all four *Erythranthe* species, the TR1 paralog was hypothesized as the ancestral copy while TR2 was the recently derived copy. The branch lengths that separated TR1 and TR2 were deep. Between *M. lewisii* TR1 and TR2 there were 241 basepairs aligned with 31 nucleotide differences, while between *M. cardinalis* TR1 and TR2 there were 251 basepairs aligned with 33 nucleotide differences. We estimated the time to the most recent common ancestor (TMRCA) between TR1 and TR2 to infer the evolutionary timing of the duplication. Assuming the genetic differences between the paralogs are neutral and a generation time of 1 per year with a mutation rate estimate from *Arabidopsis* (Ossowski et al. 2010), we estimated the duplication time at 9.19–9.39 million years ago. The timing is older than the species divergence time between *M. cardinalis* and *M. lewisii* (5.5 million years ago) (Kumar et al. 2017), suggesting the paralogs are an evolutionary old duplication that has maintained throughout the speciation of the two species.

### The conserved regions within the telomerase RNA gene reveal evolutionary changes in the templating domain

We used the functionally validated *A. thaliana* CRs (Song et al. 2019) as the reference sequence and annotated the five CR domains across the *Mimulus* TR multisequence alignment (Supplementary Fig. S5). CR2 and CR3 forms the T-PK domain with the CR1 templating domain (Mitchell and Collins 2000; Song et al. 2019) and this region in *Mimulus* had numerous indel and genetic variation between species that it was difficult to demarcate the CR2 and CR3 domain boundaries. Most noticeable was the insertions of ∼16 bp nucleotides in the *Simiolus* section species and the 50 bp insertion in the TR2 duplicate in *M. aurantiacus* ssp. puniceus reference genome. CR4 and CR5 folds into the stem-loop helix IV domain (Chen et al. 2002; Song et al. 2019) and these regions were largely conserved across *Mimulus* species.

We then focused our analysis on the CR1 domain as it holds the templating sequence and determines the telomere sequence (Fig. 1). Initially we focused on the CR1 sequences from *M. aurantiacus ssp. puniceus*, *M. guttatus*, and *M. verbenaceus* since for the three species we have determined their potential telomere sequence through genomic analysis (Fig 2 and Supplementary Fig. S2), and we could test whether the TR templating sequence would match the predicted telomere sequence motif. Results showed for each species a perfect match between the telomere sequence motif (determined through k-mer and reference genome analysis) and the TR templating sequence. For instance, *M. aurantiacus* ssp. puniceus templating sequence was AACCCTA (Fig. 3C) and the telomere sequence was determined as (AAACCCT)*_n_* (Fig. 2) [note a tandem repeat of AACCCTA sequence would be identical to a telomere motif (AAACCCT)*_n_*]. In *M. guttatus* the templating sequence was AACCCGA (Fig. 3C) and its telomere sequence was determined as (AAACCCG)*_n_* (Fig. 2). In *M. verbenaceus* the templating sequence was AACCGA (Fig. 3C) and its telomere sequence was determined as (AAACCG)*_n_* (Fig. 2).

The TR sequence results from *M. aurantiacus* ssp. puniceus, *M. guttatus*, and *M. verbenaceus* indicated the TR templating sequence corresponded to the telomere repeat motif that was synthesized at the telomere. We examined the rest of the *Mimulus* species TR sequence that was assembled from the total RNA transcriptome sequencing and focused on the CR1 domain, which holds the sequences that are responsible for synthesizing the telomere DNA. The two major sequences within the CR1 domain are the annealing sequence and the templating sequence (Fig 1). For the annealing sequence, all *Mimulus* species had an identical AACC sequence. Given that the annealing sequence is crucial for the proper template shift and double-stranded binding between the telomere DNA and TR RNA sequence, it was expected that the annealing sequence would be conserved across all *Mimulus* species. But for the templating sequence there were nucleotide differences, specifically between *Mimulus* species from different sections. We discovered all *Diplacus* section species had the templating sequence AACCCTA, all *Simiolus* section species had the templating sequence AACCCGA, and in the *Erythranthe* section the species with the TR1 orthologs had the templating sequence AACCGA.

In summary, our results indicated the *Mimulus* genus had at least two mutational changes in the telomere sequence motif. Species of the same section had identical telomere sequence motifs, but between species of different sections there was variation in the telomere sequence motif suggesting the evolutionary changes occurred in the common ancestors of each section. Based on these evolutionary changes we propose a single-step mutational model of telomere sequence evolution to explain the genetic variation within the *Mimulus* telomere. The telomere sequence motif AAACCCT is the most commonly observed sequence in plants, hence it is likely to be the ancestral telomere sequence in the *Mimulus* genus and is carried by the *Diplacus* section species. A single nucleotide mutation then converted the ancestral sequence to AAACCCG in the common ancestor of the *Simiolus* and *Erythranthe* section. Subsequently in the *Erythranthe* section, a single nucleotide deletion resulted in the AAACCG telomere sequence. It’s possible our proposed evolutionary model of *Mimulus* telomere sequence evolution may also occur in other plant species with divergent telomere sequences. We predict these other plants could have related species in the same genus with telomere sequences that are connected through single mutational changes that would ultimately trace back to the ancestral AAACCCT telomere sequence. Whether such a single-step mutational model can explain the evolutionary origins of complex telomere sequences, for instance observed in *Allium* (CTCGGTTATGGG)*_n_* (Fajkus et al. 2016), remains to be examined.

### *M. cardinalis* and *M. lewisii* TR paralogs have genetic variation in the templating sequence

We then examined the CR1 domain in *Mimulus* species with TR duplications. If the templating sequence between the TR paralogs exhibits genetic variation, this could result in the species carrying a telomere with a mixed repeat motif (*i.e.* a heterogeneous sequence structure). For the TR sequences from the *M. aurantiacus* ssp. puniceus reference genome, both TR1 and TR2 paralogs had an identical templating sequence corresponding to AACCCTA, indicating *M. aurantiacus* ssp. puniceus would have a telomere with a homogeneous sequence structure, and our genomic analysis shows that was the case (Fig. 2 and Supplementary Fig. S2). But in *M. cardinalis* and *M. lewisii* the two TR paralogs had variation within the templating sequence. The TR1 paralog had the templating sequence AACCGA, meanwhile the TR2 paralog had the templating sequence AACCCGA. This indicated *M. cardinalis* and *M. lewisii* would synthesize a telomere with a heterogeneous sequence that comprises both of AAACCG and AAACCCG motifs.

We investigated the possible functionality of the *M. cardinalis* and *M. lewisii* TR duplicates by first conducting RT-PCR of the duplicate genes on three different tissues (root, mature leaf, and floral meristem). We first compared the expression profiles of the TR gene in *M. cardinalis* and *M. lewisii*, which have the TR duplications to *M. parishii* and *M. verbenaceus*, which are the species from the same *Erythranthe* section but without the TR duplication. The *M. verbenaceus* TR expression profile and telomere sequence motif analysis were shown in Fig. 2, Fig 3, and Supplementary Fig. S2; and *M. parishii* had identical results to *M. verbenaceus*. In *M. parishii* we detected the TR gene was expressed in all of the tissue we examined (Fig. 4 bottom and see Supplementary Fig. S6 for negative PCR results when total RNA was used for RT-PCR). The TR templating sequence consisted of AACCGA (Fig. 3C) and analyzing the chromosome-level genome assembly the telomere sequence at chromosome ends was predicted to be AAACCG (Fig. 4 top). Hence, *M. parishii* and *M. verbenaceus* were synthesizing a single motif telomere sequence (AAACCG)*_n_* from its single copy TR gene.

**Figure 4.**
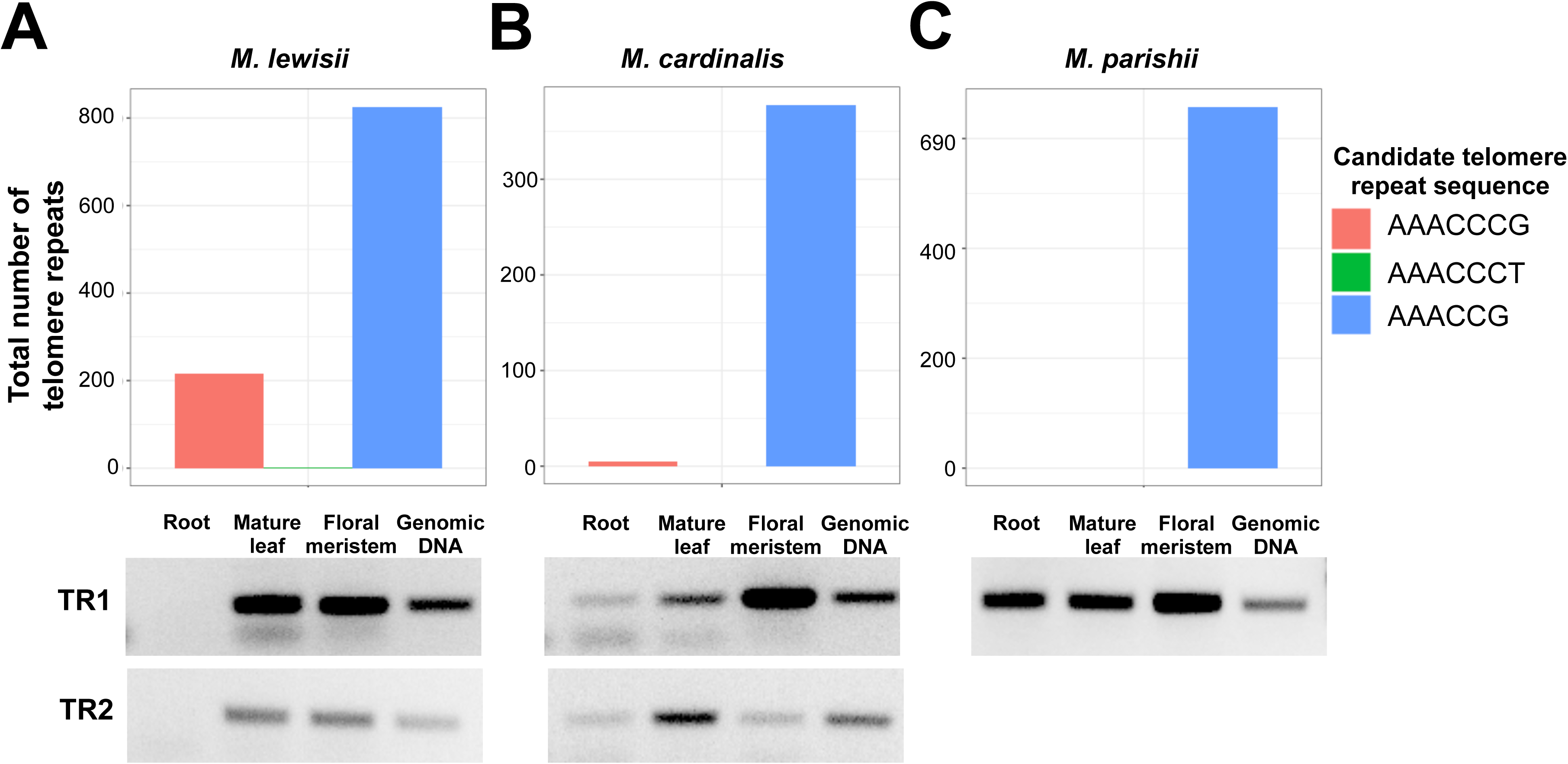
*Erythranthe* section Telomerase RNA (TR) gene duplication and its effect on the telomere sequence. Shown are telomere sequence abundance at reference genome assembly chromosome ends (above) and RT-PCR results (below) amplifying TR transcripts on cDNA generated from RNA extractions of three tissues (root, mature leaf, and floral meristem) with genomic DNA is shown as a positive control for (A) *M. lewisii*, (B) *M. cardinalis*, and (C) *M. parishii.* See Supplementary Fig. S6 for RT-PCR results from total RNA. Note *M. verbenaceus* is also a species from the *Erythranthe* section and its results have already been shown in Fig. 3A and Supplementary Fig. S2, and largely corroborated *M. parishii* results.

For *M. cardinalis* and *M. lewisii*, both TR1 and TR2 were expressed in multiple tissues (Fig. 4 bottom), especially with a notable expression in the floral meristem. Meanwhile *M. lewisii* did not have any expression of either TR gene in the root, but low expression of both TR paralogs were detected in the *M. cardinalis* root tissue. The similar TR gene expression profiles between *M. cardinalis* and *M. lewisii* suggested both species should have a telomere with a heterogeneous sequence structure. We investigated this possibility by quantifying the telomere repeat motifs at chromosome ends of *M. cardinalis* and *M. lewisii* chromosome-level reference genome assemblies, and discovered the two species had contrasting telomere sequences. In *M. cardinalis* there was only the AAACCG sequence found at chromosomal ends, meanwhile in *M. lewisii* both AAACCG and AAACCCG sequences were found at chromosomal ends (Fig. 4 top). This would occur if the expressed TR2 gene in *M. cardinalis* was not incorporated into the telomerase during the telomere DNA maintenance and only the TR1 gene was being used, suggesting the TR2 gene in *M. cardinalis* was functionally ineffectual.

### The *Erythranthe* section TR paralogs arose from an evolutionarily old gene duplication event

Our results have shown that within the *Erythranthe* section, *M. cardinalis*, *M. parishii*, and *M. verbenaceus* had the same telomere sequence structure where it consisted of AAACCG motif, while *M. lewisii* had a mix of AAACCG and AAACCCG motifs. This molecular phenotype is consistent with the underlying phylogenetic relationships where *M. cardinalis*, *M. parishii*, and *M. verbenaceus* are a monophyletic group while *M. lewisii* is a phylogenetic sister outgroup (Vickery and Wullstein 1987; Beardsley et al. 2003; Nesom 2014; Nelson et al. 2021). The TR duplication status, however, contrasted the underlying evolutionary relationship since *M. cardinalis* and *M. lewisii* shared the TR duplication but not with *M. parishii* or *M. verbenaceus*. We conducted a genomic analysis on the TR2 gene region in *M. cardinalis* and *M. lewisii*, and examined the syntenic region in *M. parishii* and *M. verbenaceus*. We searched for a TR gene within the syntenic region, which was determined through examining the orthology of the genes surrounding the TR2 gene. Results showed clear synteny across the 4 species in the genomic region surrounding the *M. cardinalis* and *M. lewisii* TR2 gene in *M. parishii* and *M. verbenaceus*, which was located at a genomic region of chromosome 7 for all 4 species (Supplementary Fig. S7). We conducted nucleotide blast analysis using the TR2 sequence as the query and searched the TR2 syntenic regions in *M. parishii* and *M. verbenaceus*. In *M. parishii* we discovered a 36 bp hit (97% identity) and in *M. verbenaceus* a 80 bp hit (88% identity), but no match that spanned the large majority of the TR2 gene was observed. These results indicated *M. parishii* and *M. verbenaceus* did not have a functional TR2 paralog and the sequences that remain in the genome are potentially pseudogenized gene remnants.

To gain a deeper understanding of the evolutionary history underlying the *Erythranthe* section TR gene duplication, we analyzed the draft genome assemblies (Liang et al. 2023) from two closely related outgroup species *M. bicolor* and *M. primuloides*. Evolutionarily, *M. bicolor* (section *Monimanthe*) is a phylogenetic sister to the *Erythranthe* section, while *M. primuloides* (section *Monantha*) is a third outgroup. The genomes for these two species were generated from short read (150 bp) Illumina sequencing and their assembly is fragmented. But through blast search we were able to discover the TR gene in both species and together with the *Erythranthe* section TR genes we reconstructed a phylogenetic tree (Fig. 5). *M. bicolor* had three TR gene copies and one paralog grouped with the *Erythranthe* section TR1 paralogs, hence we designated that copy as the *M. bicolor* TR1 gene. In addition, there were two *M. bicolor* TR paralogs that grouped with the *M. cardinalis* and *M. lewisii* TR2 paralogs, hence they were designated as *M. bicolor* TR2-1 and TR2-2. In the outgroup *M. primuloides* there was only one TR gene and phylogenetically it clustered with the TR1 paralogs, however, the bootstrap support was relatively low (bootstrap score of 70).

**Figure 5.**
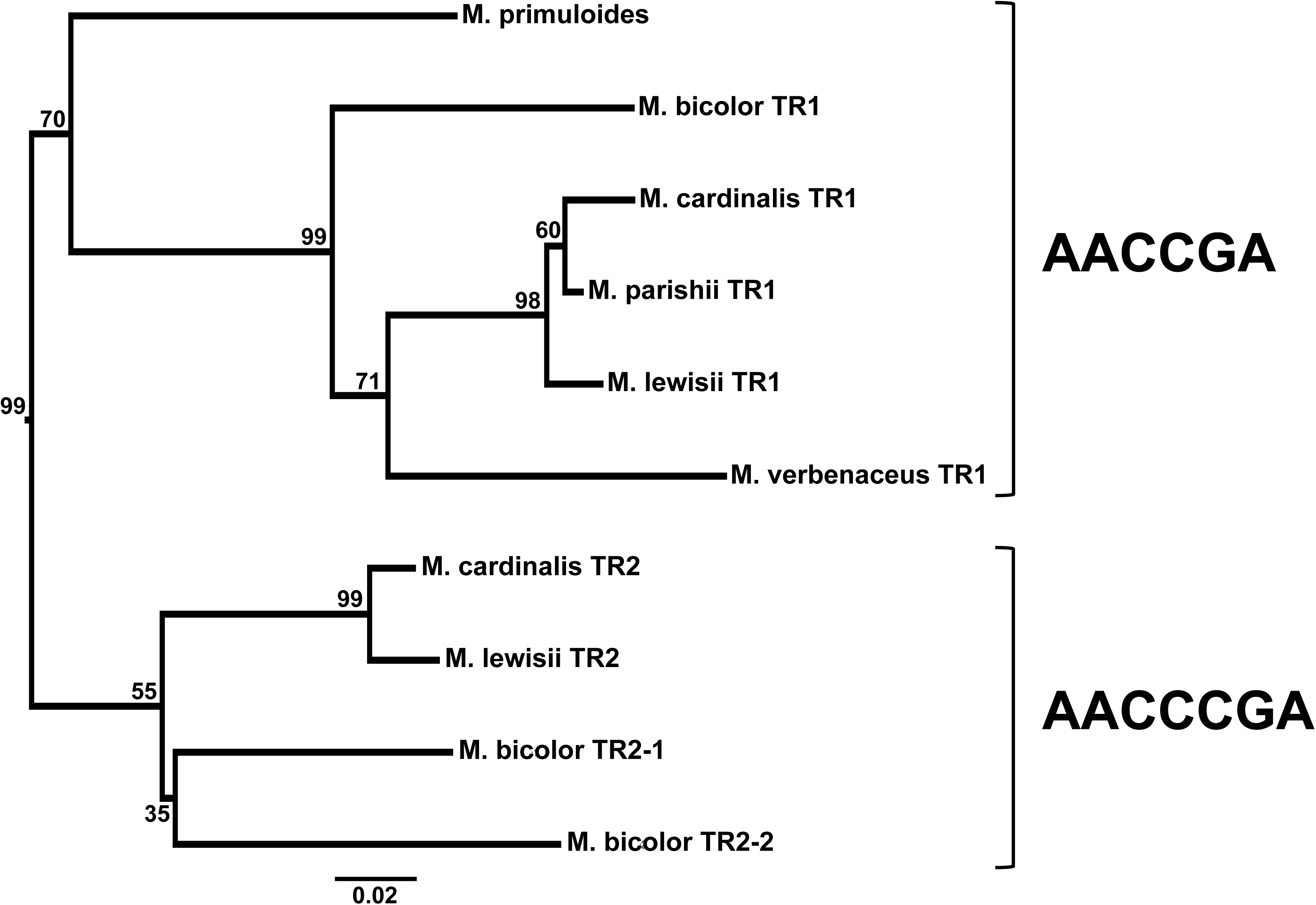
Phylogeny of the TR gene from species in the *Erythranthe* section (*M. cardinalis*, *M. lewisii*, *M. parishii*, and *M. verbenaceus*) and outgroup *M. bicolor* and *M. primuloides*. The TR gene phylogeny shows sequences grouping by paralog. The templating sequence within the TR paralog is shown on the right. Internal nodes represent bootstrap support after 1,000 replicates. A phylogeny was constructed using the sequence dataset from Fig. 3B and rooted using the *Diplacus* section (Supplementary Fig. 8), but the tree was pruned to only show the *Erythranthe* section subtree.

We examined the templating sequence of the TR genes from *M. bicolor* and *M. primuloides* to understand the evolutionary origin of the TR duplication and the resulting telomere sequence evolutionary history. We assumed an identical sequence in the templating sequence is evidence of a synapomorphic trait (*i.e.* identity inherited from the common ancestor). Results showed the TR1 monophyletic group including the *M. primuloides* TR had the templating sequence AACCGA, meanwhile the TR2 monophyletic group had the templating sequence AACCCGA. Since the outgroup *M. primuloides* had a templating sequence that was identical to the TR1 paralogs, this indicated the TR1 gene was likely to be the ancestral copy and the TR2 gene is the duplicated paralog. This also suggests the (AAACCG)_n_ is the ancestral telomere sequence for the *Erythranthe* section.

Our analysis of the *Erythranthe* section TR genes revealed an intriguing evolutionary history underlying the TR duplication. We discovered *M. cardinalis* and *M. lewisii* shared a TR duplication with an outgroup *M. bicolor* but not with the evolutionarily distant outgroup *M. primuloides*, indicating the timing of TR duplication is old and had occurred during the common ancestor of the *Erythranthe* and *Monimanthe* section. Divergence time between the two sections is estimated at ∼8.2 million years ago (Kumar et al. 2017) and our gene paralog divergence time estimate was ∼9.3 million years ago, indicating the duplication event was indeed an ancient event. After the duplication, the TR1 gene group was evolutionarily retained and all species that were examined in this study carried orthologs of the TR1 gene, suggesting the TR1 paralog is the ancestral copy. But for the TR2 gene, which we hypothesized as the recent duplicate, the paralog was lost in *M. parishii* and *M. verbenaceus*, while in *M. cardinalis* the paralog was not functionally involved in synthesizing the telomere sequence (Fig. 4). The evidence indicates the TR2 gene is pseudogenized (or in the case of *M. cardinalis* is undergoing pseudogenization) and the three species (*M. cardinalis*, *M. parishii*, and *M. verbenaceus*) are reverting to a single TR copy. For *M. lewisii,* our analysis suggests the species have retained the duplicate TR copies for a significant evolutionary time period and is functionally involved in synthesizing the telomere sequence. The possible functionality of the duplicate TR gene is currently unknown and it is intriguing that two duplicate copies of a gene with a crucial telomere maintenance function were retained for a long period. It’s possible the paralog may have acquired novel functions unrelated to telomere biology in *M. lewisii*. For instance, the *A. thaliana* TR gene was originally discovered as a long noncoding RNA gene that was differentially expressed during stress response (Wu et al. 2012), hence TR2 may have specialized stress related functions that is not observed in the TR1 paralog. Further functional experiments will be necessary to answer this question.

### The telomerase RNA gene synteny is not maintained between *Mimulus* species from different sections

During the analysis of the *M. lewisii* TR gene duplication we noticed the TR paralogs were located at two different chromosomes. In *M. lewisii*, the TR1 gene was located at chromosome 6 while the TR2 gene was located at chromosome 7 (Table 1). It was intriguing that the two duplicates were located at physically distant genomic positions and we investigated the evolutionary mechanism that led to this duplication. We first attempted to elucidate the evolutionary basis of the duplication by differentiating the TR paralogs into the ancestral versus derived form and use this information to study the evolutionary origin using synteny analysis. We hypothesized the TR1 gene was the ancestral copy (Fig. 3B and Fig. 5) and the syntenic region of the *M. lewisii* TR1 gene in *M. cardinalis*, *M. parishii*, and *M. verbenaceus* also had a TR gene located within the region (Supplementary Fig. S7 and Table 1), indicating the TR1 paralog was likely to be the ancestral TR gene within the *Erythranthe* section. We note TR1 in *M. cardinalis* is located at chromosome 7, which seemingly contradicts with the TR1 position of chromosome 6 in *M. lewisii*, but this is due to the TR1 gene located within a natural chromosomal translocation that occurred between chromosome 6 and 7, and this structural variation differs between *M. lewisii* and *M. cardinalis* (Fishman et al. 2013). In other words the TR1 gene located at two different chromosomal positions between *M. lewisii* and *M. cardinalis* are orthologs that have arisen from a shared common ancestor, meanwhile the *M. lewisii* TR1 and TR2 that are also located at two different chromosomal positions are paralogs and have arisen through a different evolutionary mechanism.

To understand the evolutionary mechanisms of the *M. lewisii* TR duplication we investigated the synteny surrounding the TR paralogs by comparing it to the evolutionarily divergent *M. guttatus* and *M. aurantiacus* ssp. puniceus reference genomes. This was conducted to examine if the TR paralog had origins from a divergent species. For instance, the templating sequence of *M. lewisii* TR2 was AACCCGA, which is identical to the templating sequence from the *M. guttatus*. We examined the *M. lewisii* TR2 gene, which was hypothesized to be the recently duplicated paralog, and investigated the synteny of the TR2 chromosomal region in *M. guttatus* and *M. aurantiacus* ssp. puniceus (Table 1). Results showed within the syntenic region there was no TR gene in *M. guttatus* (*e.g.* compare the *M. guttatus* TR gene position in Table 1 row 3 column 3 with *M. guttatus* syntenic location to *M. lewisii* TR2 gene in Table 1 row 11 column 4) and *M. aurantiacus* ssp. puniceus (*e.g.* compare the *M. aurantiacus* ssp. puniceus TR gene position in Table 1 row 4 & 5 column 3 with *M. aurantiacus* ssp. puniceus syntenic location to *M. lewisii* TR2 gene in Table 1 row 11 column 5), suggesting the *Erythranthe* section TR2 paralogs did not originate from *M. guttatus* or *M. aurantiacus* ssp. puniceus. A possible scenario to explain the origin of the TR2 paralog is if the gene arose through a transposition-mediated mechanism of gene duplication (Freeling 2009). In this case, an ancestral TR gene had been duplicated and inserted into a new chromosomal position that any evidence of common ancestry (*i.e.* synteny) had become lost. We next investigated the synteny of the *M. lewisii* TR1 gene in the evolutionarily divergent *M. guttatus* and *M. aurantiacus* ssp. puniceus reference genomes, and discovered those two species also did not have a TR gene in the syntenic region (Table 1). In other words, similar to the TR2 gene evolutionary history, the *M. lewisii* TR1 gene had also transposed and inserted into a new chromosomal position compared to *M. guttatus* and *M. aurantiacus* ssp. puniceus.

The lack of TR synteny was not limited to *M. lewisii* and was observed in all *Erythranthe* species (Table 1, see each species from the *Erythranthe* section and their TR syntenic position in non-*Erythranthe* section species that has a lack of a TR gene). For example, in *M. verbenaceus* the TR is located at chromosome 7 and when we examined the syntenic region in *M. aurantiacus* ssp. puniceus and *M. guttatus* (Fig. 6), the coding sequences surrounding the *M. verbenaceus* TR gene were clearly syntenic across the three species but there was no TR gene within the syntenic positions for *M. aurantiacus* ssp. puniceus and *M. guttatus*. The same results were observed when *M. aurantiacus* ssp. puniceus or *M. guttatus* was the focal species and the synteny of their TR gene was examined (Fig 6). In summary this indicated that species in the same section shared TR synteny, but between species of different sections the TR was located at non-syntenic chromosomal positions (Table 1). These results strongly indicated that the TR gene had frequently transposed and relocated to a different chromosomal region during the evolution of the *Mimulus* genus. We hypothesize this transposition event had occurred in the common ancestor of each section, explaining why species from different *Mimulus* sections did not share the TR gene synteny but species in the same section, for example the *Erythranthe* section, all have TRs with shared synteny. The molecular mechanism that led to the transposition of the TR gene is not known, but we speculate transposable elements may have been involved. In six of the nine *Mimulus* species where we annotated the TR gene in the reference genome, there was a transposable element sequence within 2,000 bp of the TR gene annotation and all six of them were a DNA transposon (Supplementary Table 3). In plants, DNA transposons have been hypothesized as a major mediator of gene duplication through direct or indirect consequences from the transposition mechanism of the transposon (Cerbin and Jiang 2018). We posit the transposition and duplication of the TR gene are potentially mediated through DNA transposons.

**Figure 6.**
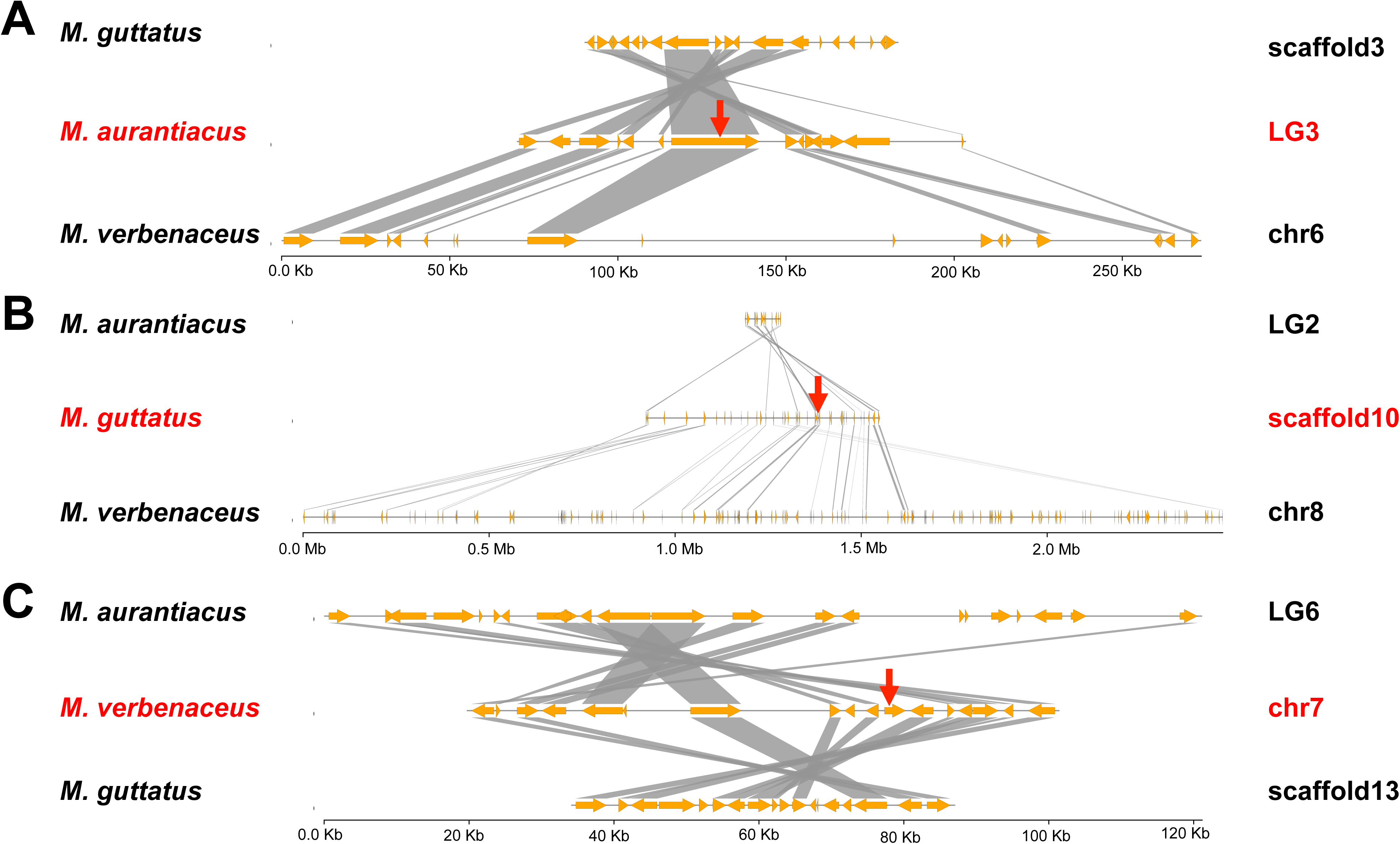
Synteny plot surrounding the Telomerase RNA (TR) gene. Each panel is showing a focal species (A) *M. aurantiacus* ssp. puniceus, (B) *M. guttatus*, and (C) *M. verbenaceus*, and its TR chromosomal location (indicated with a red arrow) is plotted in the central position and highlighted with red letters. Orange arrows indicate genes and gray boxes indicate orthology between genes. For each focal species TR gene, we show the five genes upstream and downstream with an orthology. In each plot there is clear synteny between species surrounding a focal species TR gene, but in the non-focal species there is no TR gene and instead it is located at a different chromosome (compare the TR locations in panel A, B, and C).

## Conclusion

In *Mimulus* the telomere sequence between species has changed at least twice from the ancestral AAACCCT telomere repeat sequence. The evolutionary basis of the telomere sequence variation can be explained by the genetic changes within the templating sequence of each species TR gene. We also discovered the TR gene itself had an intriguing evolutionary history within the *Mimulus* genus that involves duplication and transposing of the gene. From our results we propose the telomerase RNA transposition, duplication, and divergence (TR-TDD) model (see Fig 7 for detail) as the evolutionary mechanism that underlies the telomere sequence variation in *Mimulus*. At its core we hypothesize the TR gene can insert into new chromosomal locations through a transposition-mediated duplication mechanism and this has occurred multiple times during the evolution of the *Mimulus* genus. This duplication opens up the opportunity for sequence mutations to accumulate between the paralogs and any mutation on the TR templating sequence would lead to a species with a heterogeneous telomere sequence structure. For instance, we discovered *M. lewisii* as an example species where its telomere sequence was a heterogeneous mixture of AAACCG and AAACCCG repeats. We predict a loss of a TR paralog would revert the species back to a telomere with a homogeneous sequence structure and depending on the templating sequence of the TR paralog that wasn’t lost it may or may not lead to a sequence turnover. For instance, our results suggest for the species *M. cardinalis*, *M. parishii*, and *M. verbenaceus*, the duplicate TR2 paralog is functionally or genetically a pseudogene and unlike its sister species *M. lewisii* their telomeres consists of a single repeat sequence (*i.e.* AAACCG). Our proposed model based on *Mimulus* results may apply to other plant species as well. For example, very recently, Závodník et al. (2023) reported several plant species unrelated to the *Mimulus* genus also with evidence of the TR duplication, indicating the evolution of TR paralogs might be a common plant evolutionary phenomenon and our proposed TR-TDD model may apply widely across the plant kingdom.

**Figure 7.**
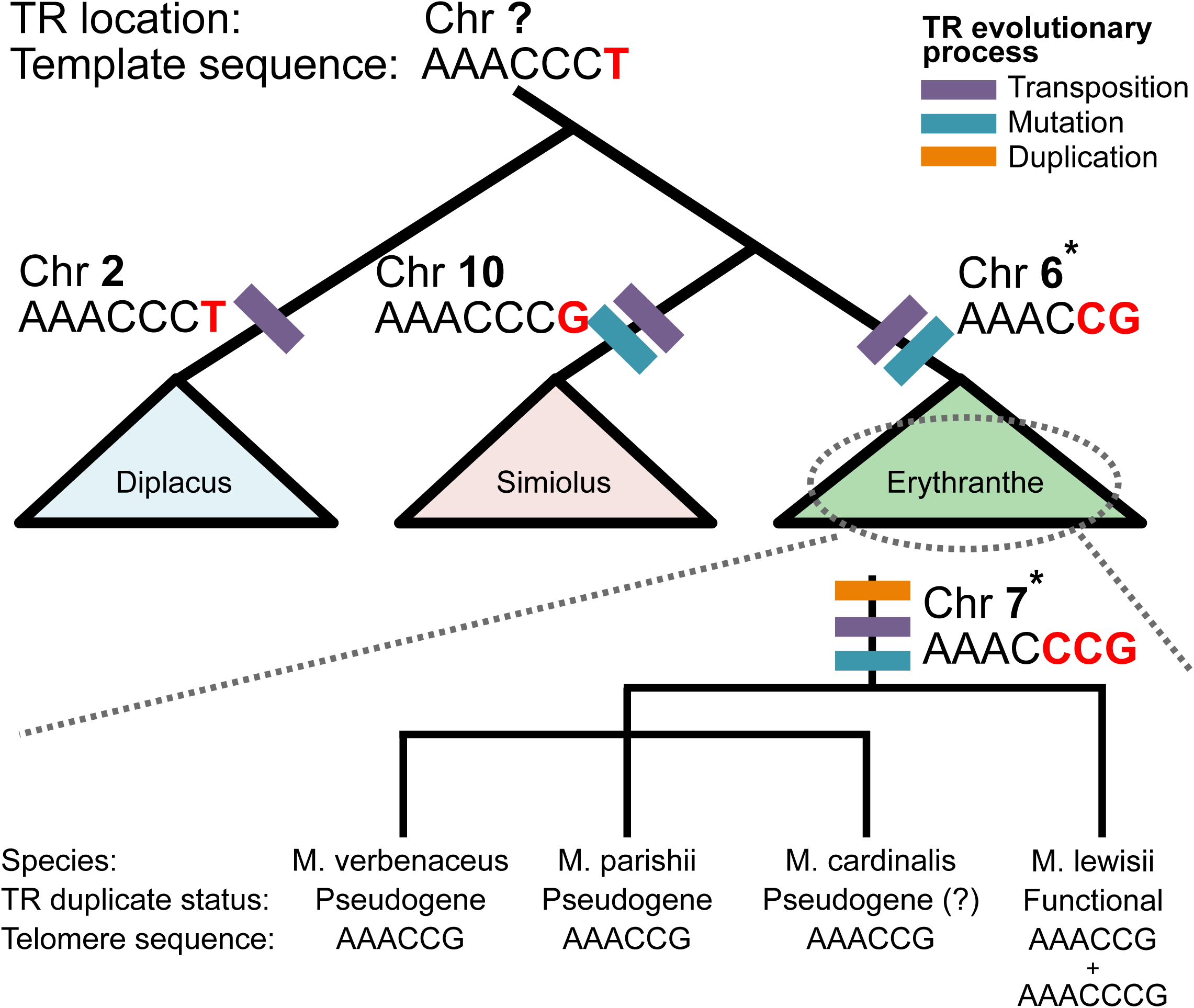
The proposed Telomerase RNA (TR) transposition, duplication, and divergence model to explain the evolutionary changes in the *Mimulus* telomere sequence. Shown are the phylogenetic relationships among the three *Mimulus* sections (*Diplacus*, *Erythranthe*, and *Simiolus*) that were examined in this study. In the branch leading up to each *Mimulus* section the evolutionary events that shaped the TR gene in the common ancestor of the section are shown with colored bars. In each branch we also show the chromosomal position of the TR gene and the TR templating sequence that ultimately determines the telomere sequence. In our evolutionary model we assume the ancestral telomere sequence in *Mimulus* to be AAACCCT and the evolutionary changes observed within the telomere sequence across the *Mimulus* genus are highlighted in red. We hypothesize the ancestral TR gene was located at an unknown chromosomal position that transposed and inserted into a new chromosomal position during the evolutionary split of each *Mimulus* section. In the *Diplacus* section the ancestral TR transposed to chromosome 2 and there was no mutation in the TR templating sequence, hence all *Diplacus* section species have the ancestral telomere sequence. Within the *Simiolus* and *Erythranthe* section in addition to the section-specific TR transposition events, the TR templating sequence was mutated resulting in species in those sections to have a telomere sequence that is AAACCCG and AAACCG respectively. In the *Erythranthe* section (magnified below) there was TR gene duplication that occurred in the common ancestor of the *Erythranthe* and *Monimanthe* section, resulting in the TR1 and TR2 gene duplicates. We posit TR2 as the recently derived paralog and it originated from a transposition-mediated gene duplication process. The two TR paralogs encode different templating sequences and *M. lewisii* is utilizing both paralogs during telomere synthesis, resulting in its telomere with a sequence heterogeneous structure consisting of both AAACCG and AAACCCG repeats. In *M. parishii* and *M. verbenaceus* the TR2 duplicate has pseudogenized, meanwhile in *M. cardinalis* it might be undergoing pseudogenization, hence in all three species the telomere consists of a single telomere repeat sequence AAACCG. * denote the chromosomal position of the TR based on *M. lewsii* genome coordinates.

There is an important question that arises from our results; what benefit, if any, would duplicate copies of the TR gene have for the plant? Currently, we do not have the functional explanation for the TR duplication. Most gene duplicates become nonfunctional (Force et al. 1999), unless it has functional consequences that lead to its retention and maintenance through selection (Kuzmin et al. 2022). A variety of models have been proposed to explain the evolutionary retention of gene duplicates (Innan and Kondrashov 2010) and may apply to the TR duplicates. For instance, the TR paralog may have gained novel function or partially redundant telomere-related function through neofunctionalization or subfunctionalization processes (Ohno 1970). It’s also possible the duplication results in changes in the gene expression of the TR and if an increase in the gene dosage is beneficial then the gene duplication may become fixed in a population through positive selection (Kondrashov et al. 2002). Understanding the functional consequences arising from the TR duplication is an important follow up research direction from our results and would lead to a deeper understanding on why the telomere sequence has changed across multiple lineages across the plant kingdom.

## Materials and Methods

### Analyzed reference genomes

All *Mimulus* reference genomes were downloaded from Mimubase (http://mimubase.org/). Specifically, the analyzed genome versions were *M. guttatus* v2.0 (Hellsten et al. 2013), *M. aurantiacus* ssp. puniceus v1.0 (Stankowski et al. 2019), *M. cardinalis* v2.0, *M. lewisii* v2.0, *M. parishii* v2.0, and *M. verbenaceus* v2.0. We note the genome sequences for *M. cardinalis*, *M. lewisii*, *M. parishii*, and *M. verbenaceus* are early access publicly available versions and a manuscript describing the biological properties of those genomes will be published elsewhere.

### k-Seek based telomere sequence analysis

We downloaded *M. aurantiacus*, *M. guttatus*, and *M. verbenaceus* whole genome resequencing data (Troth et al. 2018; Stankowski et al. 2019; LaFountain et al. 2023) from NCBI SRA with the identifiers PRJNA549183, PRJNA344904, and PRJNA813304. Raw reads were trimmed using BBTools (https://jgi.doe.gov/data-and-tools/bbtools/) bbduk.sh script version 39.01 with parameters minlen=25 qtrim=rl trimq=10 ktrim=r k=25 mink=11 hdist=1 tpe tbo. Trimmed reads were then analyzed with k-Seek (Wei et al. 2014; Wei et al. 2018) to quantify the total copy number of a tandem repeat sequence. k-Seek requires a k-mer repeat length to be at least 50 bp in order to avoid counting small tandem repeats scattered across the genome. To account for the differences in genome coverage between samples we normalized each sample’s k-mer counts by the sample’s median genome coverage. Genome coverage was calculated by first aligning the whole genome resequencing reads to the reference genome of the respective species using bwa-mem version 0.7.16a-r1181 (Li 2013) and then the per-sample average coverage was calculated using bedtools version 2.25.0 (Quinlan and Hall 2010).

### Reference genome based telomere sequence analysis

All *Mimulus* species have a reference genome that is assembled into chromosomes (*i.e.* linkage groups), hence if the telomere sequence was assembled it would be at the ends of each chromosome assembly. From each species reference genome we extracted the starting and ending 500 bp of the assembled chromosome sequence using samtools version 1.14 (Li et al. 2009). For each extracted sequence we then searched for the telomere repeat sequences AAACCCT, AAACCCG, or AAACCG using bash command grep with option -aob and counted the number of matches. The data was visualized using ggplot2 package (Wickham 2011) in R 4.3.1 (RStudio 2020).

### Plant material

The *Mimulus* species that were used in this study with the growth conditions are mentioned in Supplementary Table 1. Total genomic DNA was isolated from the leaves of *M. cardinalis*, *M. guttatus*, *M. aurantiacus* ssp. puniceus, *M.parishii, M. verbenaceus*, and *M. lewisii* using the modified CTAB method (Doyle 1990).

### RNA isolation and reverse transcription-polymerase chain reaction (RT-PCR)

The total RNA was extracted from 100 mg of floral meristem, root, and leaves using TRIzol (Invitrogen) according to the manufacturer’s protocol. The RNA was further treated with DNaseI (RNase-free, New England Biolabs) followed by column purification using Monarch Total RNA Miniprep Kit (New England Biolabs). The ProtoScript First Strand cDNA Synthesis Kit (New England Biolabs) was used for the synthesis of cDNA using the manufacturer’s protocol. One µg of RNA was reverse transcribed using random primers.

RT-PCR was performed on the root, leaves, and floral meristem of *M. cardinalis*, *M. guttatus*, *M. aurantiacus* ssp. puniceus, *M.parishii, M. verbenaceus*, and *M. lewisii*. Four µl of 10X diluted cDNA was used in a 20 µl reaction mix. Taq DNA Polymerase with Standard Taq Buffer (New England Biolabs) was used for the amplification of TR under conditions as follows: 10 min at 95°C; 30 sec at 95°C; 30 cycles of 30 sec at 95°C, 30 sec at 55°C, 15 sec at 68°C; and final extension at 68°C for 5 min. The primers used for the RT-PCR are listed in the Supplementary Table 4. The PCR products were electrophoresed at 95 volts for 40 minutes on a 2 % agarose gel stained with ethidium bromide.

### RNA-seq library preparation

The total RNA extracted from the floral meristem of species listed in Supplementary Table 1 were checked for quality using RNA ScreenTape (Agilent Technologies) and Qubit 2.0 fluorometer was used for determining the concentration. From the total RNA the ribosomal RNA (rRNA) were depleted using QIAseq FastSelect–rRNA Plant Kit. Using rRNA depleted RNA samples the RNA-seq libraries were prepared using NEBNext Ultra II Directional RNA Library Prep Kit according to the manufacturer’s protocol. RNA-seq libraries were sequenced by the University of Kansas Genome Sequencing Core on a Illumina NextSeq 2000 platform using P3 Reagents and as 2 × 100 bp paired-end reads.

### *De novo* transcriptome assembly

From the raw sequencing reads, the quality of reads were analyzed using fastqc version 0.11.8 (Andrews 2010). The reads were subjected to quality control trimming using the bbduk.sh script in the BBMap package version 39.01 (Bushnell et al. 2017). The processed data was used for *de novo* transcriptome assembly using the Trinity pipeline version 2.8.5 (Grabherr et al. 2011) and following their assembly tutorial (https://github.com/trinityrnaseq/trinityrnaseq/wiki). The *de novo* assembly was runned as paired-end option with the command: Trinity --seqType fq --left “$FQ1” --right “$FQ2” --CPU 40 --max_memory 140G --output “$OUTDIR”. To align the reads back to the reference genome and check for read alignment statistics we used the program Bowtie2 version 2.3.5.1 (Langmead et al. 2009). TrinityStats.pl script from the Trinity pipeline was used to compute transcript contig N 50 values. To quantify the completeness of the transcriptome assemblies the Benchmarking Universal Single-Copy Orthologs (BUSCO) v3.0.2 (Simão et al. 2015) was executed with the following command: run_BUSCO.py -c 20 -o OUTPUT_NAME --in SEQUENCE_FILE -l eudicotyledons_odb10 -m transcriptome.

### Telomerase RNA gene bioinformatic analysis

Candidate TR gene annotation was conducted using a previously implemented bioinformatics methodology that was used for detecting the TR in various land plant species (Song et al. 2019). We used TR sequence from previously annotated plants to generate a position weight matrix (PWM) to search for the candidate TR sequences within the *Mimulus* reference genomes using fragrep2 (Mosig et al. 2007), and the program Infernal (Nawrocki and Eddy 2013) was used to search for candidate TRs using secondary structure and sequence similarity.

To search for the TR sequence within the *de novo* transcriptome assembly we used the TR gene that was annotated from the reference genome using fragrep2 and Infernal. Using the standalone blast+ version 2.9.0 (Camacho et al. 2009) the candidate TR gene in the transcriptome assemblies were identified using the following command: blastn -query “$QUERY_FASTA” -db “$REF_DB” -task blastn -outfmt 6. The closely related *Mimulus* species from the same section was used as the query sequence. The TR gene detected from each *Mimulus* species were aligned to each other using MUSCLE (Edgar 2004) with default parameters on MEGA (Kumar et al. 2018).

### Annotating conserved and functional regions in the *Mimulus* TR

For species with the reference genome, the 100 bp upstream region of each species TR gene was retrieved for analyzing the conserved regulatory elements. The sequences were aligned to each other using MUSCLE (Edgar 2004) with default parameters on MEGA (Kumar et al. 2018). We specifically amplified the *M. cardinalis* TR2 upstream region and Sanger sequenced using primers mentioned in Supplementary Table 4. In addition, we annotated the functional domains with the *Mimulus* TR sequence using *A. thaliana* TR sequence. Since previous *A. thaliana* study had experimentally validated the secondary structure and functional domains of the TR (Song et al. 2019), we aligned the *A. thaliana* TR sequence with the *Mimulus* TR multi-sequence alignment and annotated the *Mimulus* TR functional domains.

### Phylogenetic analysis

The multi-species TR gene alignment was used for the phylogenetic analysis with RAxML ver. 8.2.5 (Stamatakis 2014) to build a maximum likelihood-based tree. We used a general time-reversible DNA substitution model with gamma-distributed rate variation and bootstrap analysis was conducted with 1000 replicates.

Divergence time between the two TR duplicates was estimated using Nei’s genetic distance (Nei 1972) (*D = 2*□*T* where *D* is genetic divergence between the gene paralogs, □ is the mutation rate per generation, and *T is* number of generations). We used mutation rate estimate from *A. thaliana* (7E-9 base substitutions per site per generation) (Ossowski et al. 2010) for L and assumed 1 generations per year.

### Synteny analysis

Synteny of the TR gene was determined through orthology of the coding sequence surrounding the TR. For each *Mimulus* species with a reference genome we took the chromosomal position of the TR and extracted the gene annotations that were up and downstream of the TR gene. For the focal species and its TR gene, orthology of the surrounding genes were determined through Orthofinder version 2.5.5 (Emms and Kelly 2015; Emms and Kelly 2019) and using gene annotations of *Mimulus* species downloaded from Mimubase (http://mimubase.org/). In order to visualize the synteny, pyGenomeViz was used by following the tutorial (https://github.com/moshi4/pyGenomeViz) (Shimoyama 2022).

### Data availability

Raw sequencing data have been deposited on the Sequence Read Archive (SRA) under the identifier PRJNA1049363. Data that was used for downstream analysis are deposited at Zenodo (doi:10.5281/zenodo.10266656).

## Acknowledgements

We thank members of the Choi lab, Askhan Shametov and Stephanie Sage for their help in preparing this study. We also thank the University of Kansas Genome Sequencing Core for the support during the transcriptome sequencing. This work was supported by grants from the National Science Foundation under award number IOS-2204729 and the National Institute of General Medical Sciences of the National Institutes of Health under award number P20 GM103418 and P30 GM145499 to J.Y.C.

---

Supplementary Table 1. The *Mimulus* species that were studied in this research.

Supplementary Table 2. The *Mimulus* transcriptome *de novo* assembly statistics.

Supplementary Table 3. The transposable element sequence directly adjacent to the TR gene.

Supplementary Table 4. The primer sequences that were used in this research.

Supplementary Fig. S1. Genome-wide tandem repeat profiles for species (A), (B), and (C). Top 25 most abundant k-mers are shown with the k-mers ordered alphabetically then by size.

Supplementary Fig. S2. Telomere repeat sequence counts at ends of reference genome assembly.

Supplementary Fig. S3. Upstream 100 bp of sequence alignments of candidate TR genes from reference genomes.

Supplementary Fig. S4. RT-PCR results amplifying TR transcripts on cDNA generated from RNA extractions of three tissues (root, mature leaf, and floral meristem) and the raw RNA as negative control. Genomic DNA is shown as a positive control.

Supplementary Fig. S5. Alignment of the entire *Mimulus* TR gene. The functional domains CR1-CR5 are indicated above the alignment.

Supplementary Fig. S6. RT-PCR results amplifying TR transcripts on cDNA generated from RNA extractions of three tissues (root, mature leaf, and floral meristem) and the raw RNA as negative control. Genomic DNA is shown as a positive control.

Supplementary Fig. S7. Synteny plot of the *M. lewisii* and *M. cardinalis* TR1 and TR2 region in *M. parishii* and *M. verbenaceus*. The TR chromosomal location is indicated with a red arrow. Orange arrows indicate genes and gray boxes indicate orthology between genes. We show the five genes upstream and downstream with orthology.

Supplementary Fig. S8. Phylogeny of the entire TR sequences assembled from 18 *Mimulus* species/subspecies total RNA transcriptome (Fig. 3B) and *M. bicolor* and *M. primuloides* TR sequences. Nodes with bootstrap support >95% are indicated with a red circle.

