## Supplementary figures and images for "Transposition, duplication, and divergence of the telomerase RNA underlies the evolution of *Mimulus* telomeres"

### Supplementary Fig S1

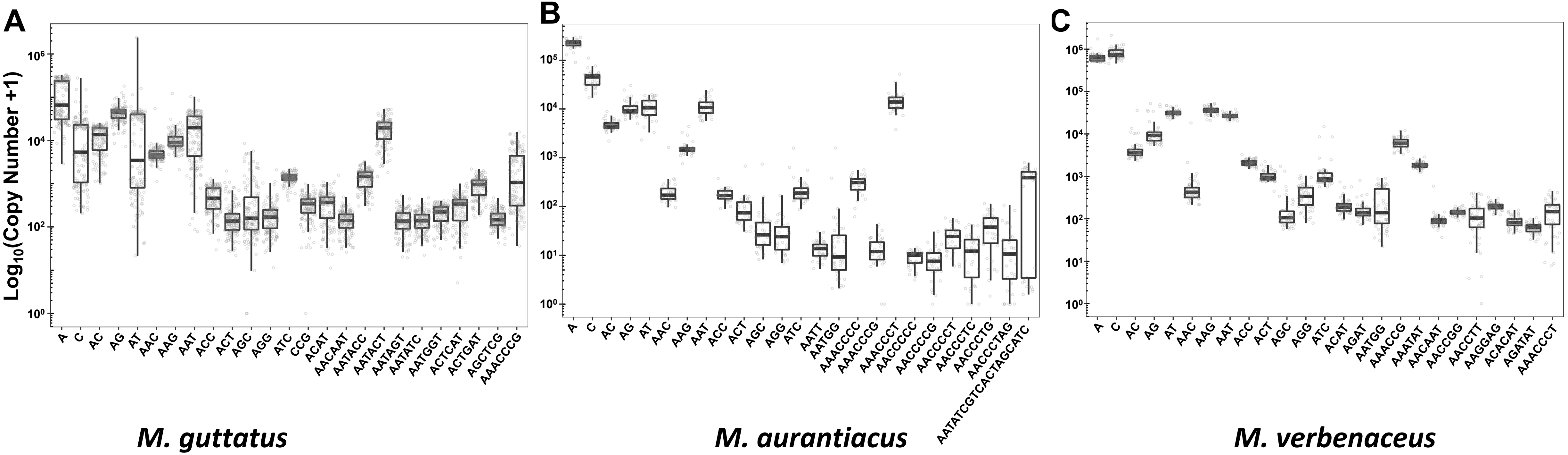

### Supplementary Fig S2

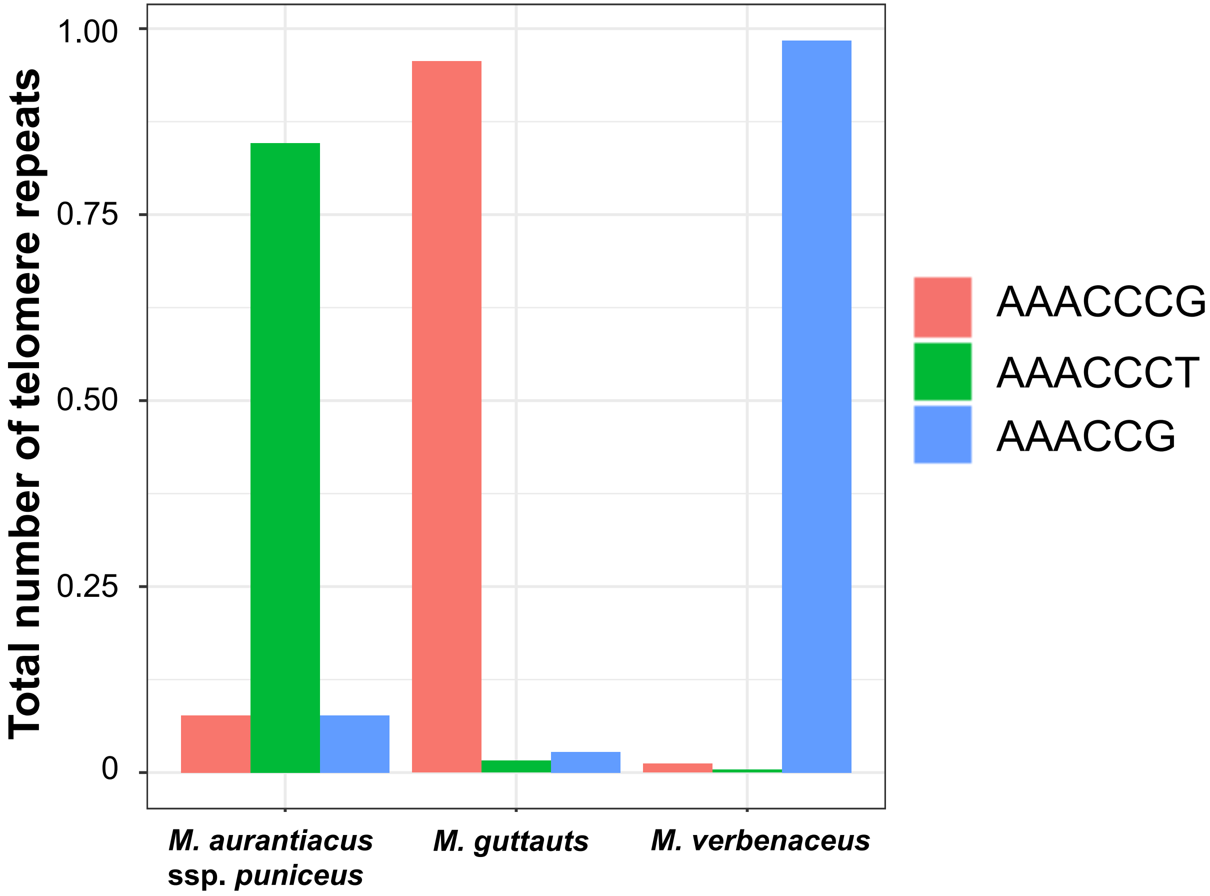

### Supplementary Fig S3

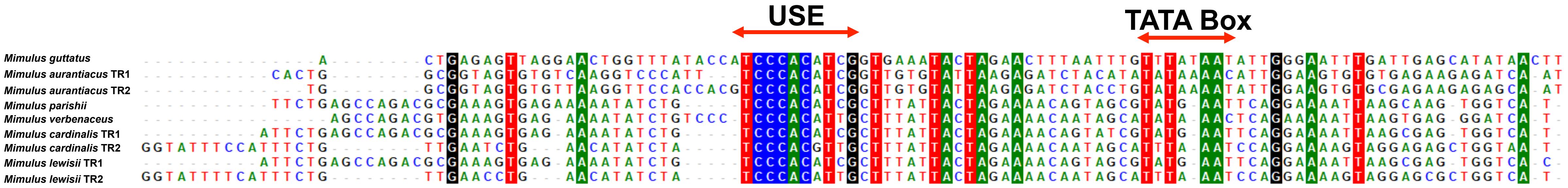

### Supplementary Fig S4

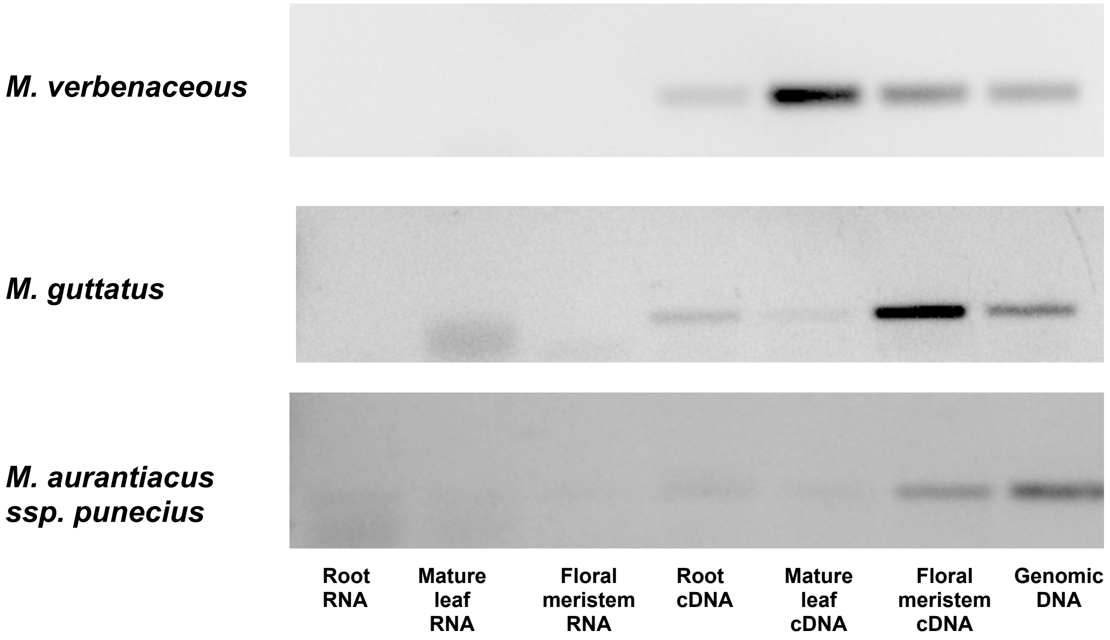

### Supplementary Fig S5

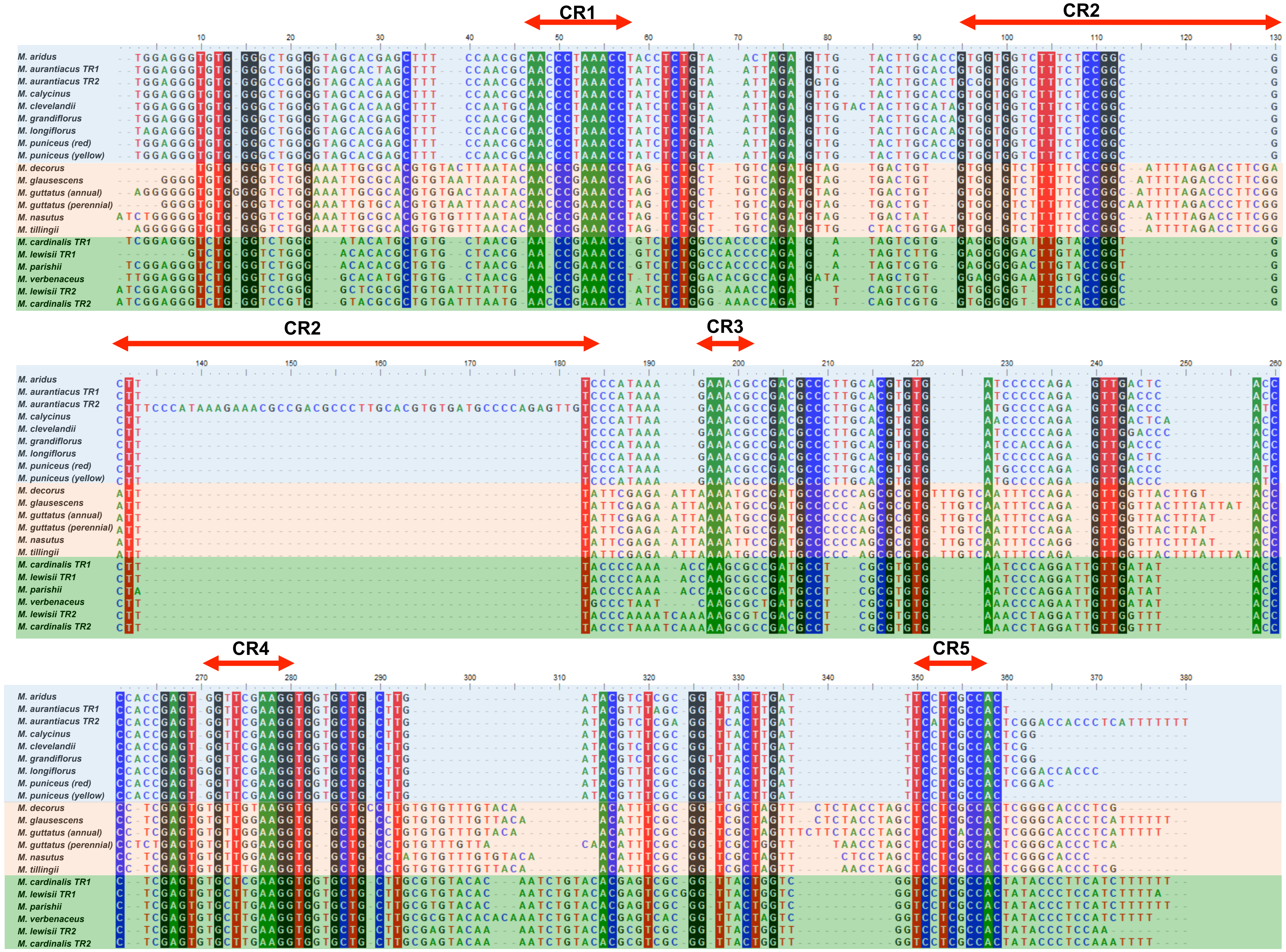

### Supplementary Fig S6

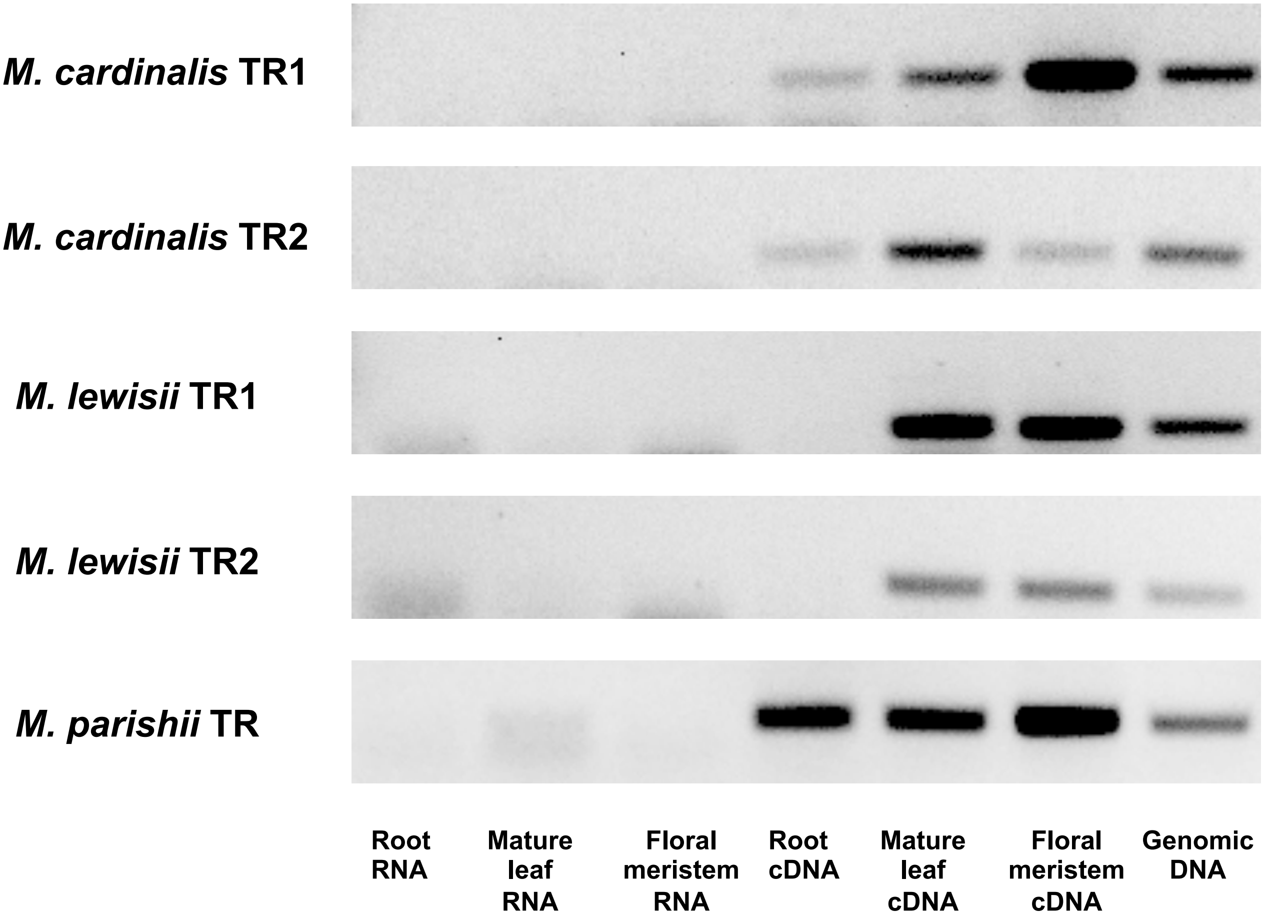

### Supplementary Fig S7

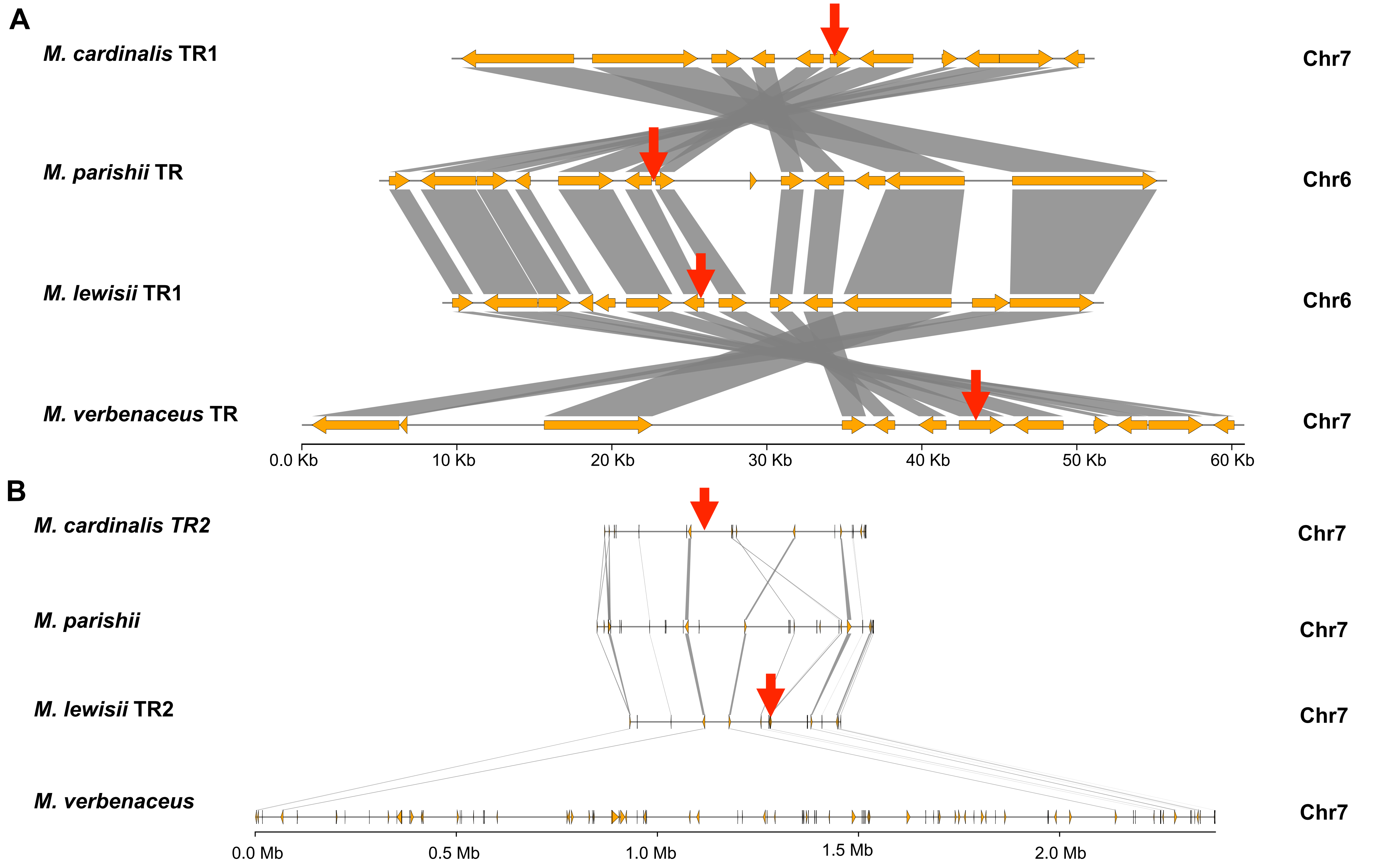

### Supplementary Fig S8

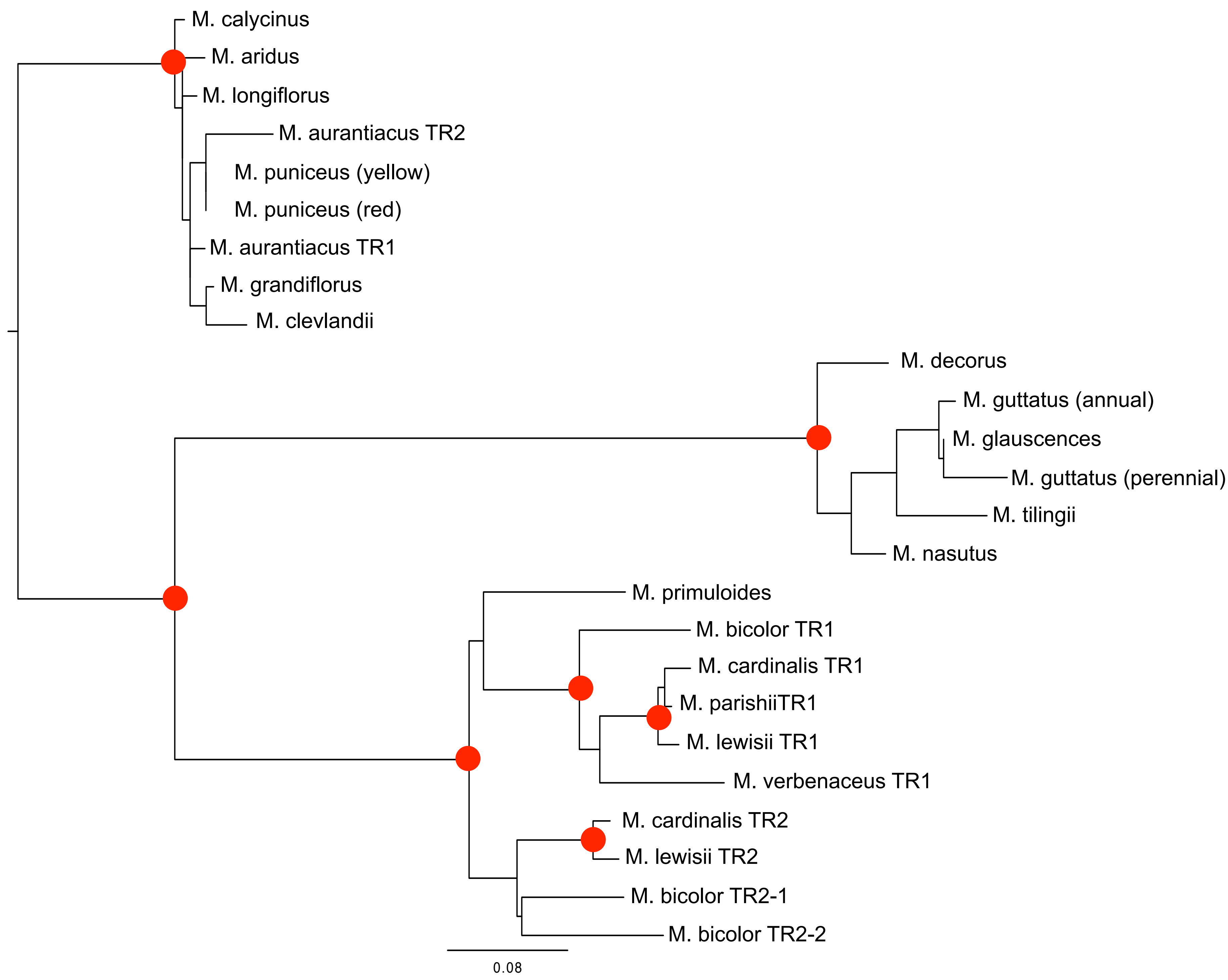
